# Structural evolution of the estrogen receptor regulatory domain

**DOI:** 10.1101/2024.12.18.629305

**Authors:** Daniel P. McDougal, Jordan L. Pederick, Linda Shearwin-Whyatt, Frank Grützner, John B. Bruning

**Affiliations:** Institute for Photonics and Advanced Sensing (IPAS), School of Biological Sciences, The University of Adelaide, Australia; School of Biological Sciences, The Robinson Research Institute, The University of Adelaide, Adelaide, Australia

## Abstract

Estrogen receptors (ERs) α, β and γ are ligand-dependent transcription factors that regulate vertebrate reproduction, cell survival and other physiological processes. Here, we report an integrative analysis of mammalian and teleost receptors, including the first structure of an ERγ ligand binding domain (LBD), showing that structural divergence acquired during evolution is accommodated by different strategies within functional regions to preserve intrinsic regulatory mechanisms. Supervised-learning and comparative Markov modelling uncover highly constrained network structures essential for allostery and protein folding, revealing a fundamental regulatory architecture and source of selective pressure in ER+ breast cancer and reproductive disorders. Finally, cross-referencing molecular constraints to genome sequencing reveal evolutionary origins underlying natural genetic variation in humans and widespread disruption of constraints. This work provides structural insights into the conservation of gene regulation by essential transcription factors and has implications for precision medicine.

## Main text

Estrogen receptors (ERs) are ligand-dependent transcription factors that regulate reproductive biology, development and cell survival, among other vital physiological processes in vertebrates^1–5^. They evolved from a common estradiol (E2)-sensitive ancestor and three ER paralogs are reported in vertebrates: ERα, ERβ and ERγ, which share structurally conserved domain architecture (**Fig. 1a** and **b**). ERα and ERβ are expressed ubiquitously (specifically in jawed-vertebrates), whereas ERγ is exclusive to teleost fish, having evolved from an ancestral ERβ during the teleost-specific whole genome duplication ^6–9^. The ERs are unusual in that regulatory functions of the ligand binding domain (LBD) have remained evolutionarily conserved despite widespread sequence variation and multiple gene duplications. This contrasts with their neighbouring 3-ketosteroid receptors (kSRs; such as the mineralocorticoid and glucocorticoid receptor), which acquired new functions following duplication and have been used as models to study functional protein evolution^7,10–12^. This dichotomy offers a unique opportunity to explore how ligand-dependent regulatory mechanisms of key vertebrate transcription factors are preserved across evolutionary time, and the molecular hurdles that must be overcome to protect gene regulation.

**Fig. 1.**
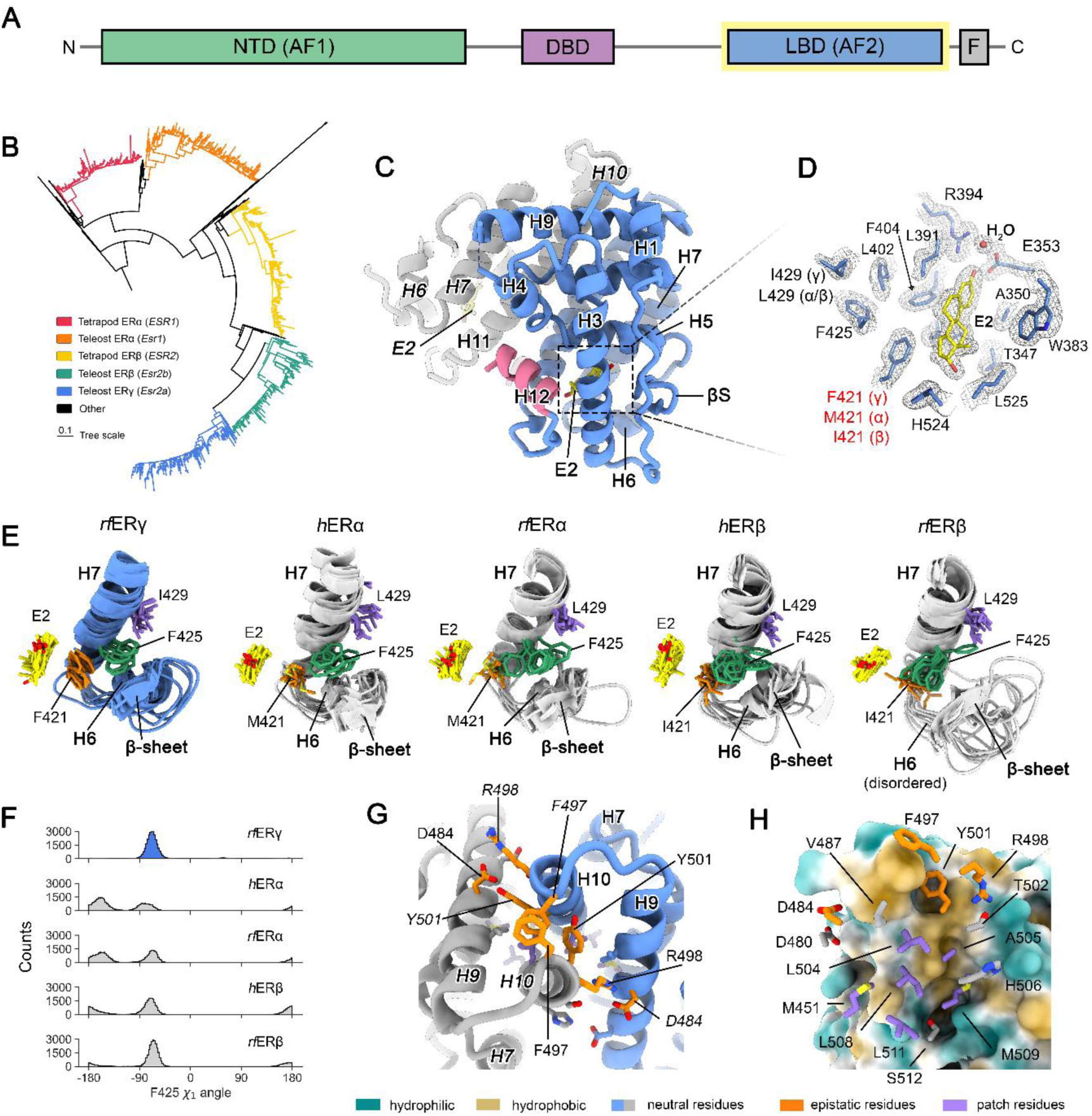
X-ray crystal structure of the rfERγ LBD reveals preservation of gene regulatory functions. **a**, Domain architecture of full-length ERs (NTD: N-terminal domain; AF1: activation-function 1; DBD: DNA-binding domain; LBD: ligand-binding domain; AF2: activation-function 2). **b**, Maximum-likelihood phylogenetic tree of 1,051 ER LBD sequences from vertebrates and protostomes; the branch corresponding to protostomes is compressed. **c**, X-ray crystal structure of the *rf*ERγ LBD solved at 1.95 Å resolution revealing conservation of protein folding, homodimer assembly, AF2 conformation and **d** E2 binding mode. Secondary structure elements are labelled in bold and residues are labelled accordingly (using *h*ERα LBD numbering for consistency). In **d**, residues that are conserved with ERα and ERβ homologs from human and rainbowfish are coloured in black. Sidechains and E2 are shown as sticks, and the simulated annealing composite omit 2F_o_-F_c_ electron density map (1.5 σ) is shown as blue mesh. **e**, Representative snapshots of the F425 χ_1_ sidechain dihedral angle obtained from K-means clustering (k=10) of triplicate 1 µs all-atom molecular dynamics (MD) simulations. Residue sidechains and E2 are shown as sticks (E2: yellow; F/M/I421: orange; F425: green; I/L429: purple). **f**, Histogram showing aggregate F425 χ_1_ sidechain dihedral angle populations for each homolog. **g**, Top-down perspective of the *rf*ERγ LBD homodimer showing structural plasticity of the dimerisation interface; notable residues are shown as sticks and coloured according to the legend below. **h**, Hydrophobic surface representation of one subunit from the *rf*ERγ LBD homodimer and opposing residue sidechains.

The ER LBD functions as a transcriptional control centre by mediating E2 binding, coactivator recruitment and homodimerisation^13,14^. It adopts an α-helical wedge fold with a central hydrophobic ligand binding pocket (LBP) that recognises E2 and assembles into a ‘head-to-head’ homodimer, for binding to DNA in the context of full-length receptor^15^. Upon E2 binding, a conformational shift occurs forming the activation function-2 (AF2) interface to where transcriptional coactivators are recruited via a conserved *LxxLL* motif^16–18^. The intramolecular pathways that allosterically couple the AF2 to ligand and distal regions of the domain during gene regulation, and the evolutionary constraints these pathways impose, are poorly understood.

Present understanding of regulatory mechanisms are derived from structural and biochemical studies of mammalian receptors^13–19^. However, despite functional homology, structural information from evolutionarily divergent vertebrates remains scarce. This gap hinders ability to address fundamental questions, such as how structural changes arising from sequence variation are tolerated so as to not disrupt regulatory functions. These principles are also important for understanding the origins of phenotypically diverse ER-related diseases, like ER+ breast cancer and reproductive disorders, which stem from mutations^20–26^. Here, we systematically address these gaps in knowledge through the characterisation of evolutionarily divergent structure, integrated with supervised-learning, Markov modelling, and analysis of human genome sequencing. This work provides a structural basis for conservation of gene regulation across vertebrates, providing key insights into the neutral evolution of an important transcription factor regulatory domain and the implications for human health.

## Results and discussion

### *X-* ray crystal structure of the ERγ LBD reveals variation within functional regions

We first aimed to determine whether E2-binding, coactivator recruitment and homodimerisation are structurally conserved between human and teleost. We solved the first crystal structure of an ERγ LBD bound to E2 at 1.95 Å resolution from the native Australian freshwater teleost fish, *Melanotaenia fluviatilis* (Murray River rainbowfish; *rf*ERs)^9,27,28^, as well as the *rf*ERα LBD bound to E2 and a human SRC2 coactivator peptide at 2.75 Å (**Table S1**). Diffracting crystals of the *rf*ERβ LBD could not be obtained, thus an energy-minimised computational model was built using the *h*ERβ LBD as a template. Notably, *h*ERs and *rf*ERs share approximately 63% pairwise sequence identity over the length of the LBD (**Extended Data Fig. 1a**). Herein we adopt *h*ERα LBD amino acid numbering for consistent interpretation. E2-binding, coactivator interaction and homodimerisation mode were found to be structurally similar between ERα and ERβ orthologs (**Extended Data Fig. 1b-d**). We then compared the *rf*ERγ LBD to human and teleost ERα and ERβ LBDs to identify changes that may have arisen during divergence of the duplicated gene. The crystal structure of the *rf*ERγ LBD exhibits conserved tertiary folding, with an average heavy-atom RMSD of 0.8 Å compared to ERα and ERβ LBDs and was captured as a ‘head-to-head’ homodimer, maintaining conservation of quaternary assembly (**Extended Data Fig. 1c**). The AF2 conformation and positioning of E2 binding mode are structurally similar with ERα and ERβ LBDs (**Fig. 1c** and **d**). This, by extension, supports the preservation of allosteric regulation. Fluorescence polarisation assays also confirmed that *rf*ER LBDs can specifically recognise the *LxxLL* motif of a non-native coactivator peptide demonstrating that the mode of coregulator recognition is conserved (**Fig. S1**). Furthermore, structural modelling of the *LxxLL* motif to *rf*ERβ and *rf*ERγ LBDs predicts the same key interactions within the AF2 (**Extended Data Fig. 1d**). These observations are in line with previous cell-based transactivation studies of teleost ER LBDs in human cell lines which reported comparable responses to E2, demonstrating the compatibility of ER LBD-mediated transcriptional machinery between mammals and teleosts^8,29,30^. Together, these data show that transcriptional regulatory mechanisms of the LBD are conserved between evolutionarily divergent vertebrates, separated by at least 420 million years^28^.

Despite the overall structural conservation of functional regions, analysis of the *rf*ERγ LBD revealed deviations within the LBP and at the dimerization interface compared to the other receptors. Within the *rf*ERγ LBP, F421 replaces M421 (ERα) or I421 (ERβ), which lies adjacent to E2 and coordinates the hormone to hydrogen bond with H524^31^. We modelled the F421 substitution and possible rotamer orientations into human and teleost ERα and ERβ backgrounds which predicted clashes with E2 and the surrounding structure (**Extended Data Fig. 2a**). However, our crystal structure shows that E2 binding is maintained for *rf*ERγ; F421 is also observed in other teleost ERγ orthologs and does not hinder specificity nor response to E2 binding^6,8,30,32^. Additionally, analysis of the dimerization interfaces shows widespread changes in surface structure resulting from sequence variation (**Extended Data Fig. 3a,b**). Unlike the interfaces of ERα and ERβ homodimers which remain mostly symmetrical, the interfaces of the *rf*ERγ homodimer are clearly asymmetric. These evolutionary changes are of particular interest because mutations which switch residue chemistry and structure within these key functional regions cause loss-of-function in *h*ERs, and geometric surface complementarity is considered, in general, essential for stable protein-protein interaction^15,25,33–35^.

### Structural plasticity preserves E2 binding in the ERγ LBP

During the evolution of a protein, the degeneracy of residue substitutions and epistasis (the non-additivity of mutational effects) can maintain neutrality of sequence variation and preserve function^36–38^. We hypothesised that degeneracy and epistasis might be important for retaining ligand-dependent regulatory mechanisms between divergent evolutionary trajectories. To address this hypothesis, the preservation of hormone binding was first investigated. In the *rf*ERγ LBD, E2 sits parallel/adjacent to the phenyl ring of F421 which clashes with the conserved F425 forcing an outward conformation nestled, between the β-sheet, H6 and H7 (**Fig. 1d** and **Extended Data Fig. 2a**). Comparison to ERα and ERβ structures shows that the substitution D/N411S relaxes the secondary structure of H6 and lengthens the β_2_-strand, while substitution of L429 for isoleucine alleviates clashes with F425. However, between ERα and ERβ structures we also found sequence-dependent changes to these structural elements (**Extended Data Fig. 2b**). To determine how the F421 substitution is tolerated and whether structure plays an important role, we performed replicate all-atom molecular dynamics (MD) simulations. In *rf*ERγ, the F425 conformation remains stable and rarely samples an alternate state. Alternatively, in ERα and ERβ homologs, F425 transitions between a spectrum of conformations. Clustering of the F425 χ_1_ sidechain dihedral angle and secondary structure analysis revealed conformational heterogeneity of surrounding structure, but that the dynamics of F425 are independent, which rotates below I/L429 (**Fig. 1e** and **f**, and **Extended Data Fig. 2c**). Importantly, the positioning of E2 is stable despite profound structural and dynamical changes between homologs. Together, these data suggest that neutrality of the F421 substitution arises from an intrinsic structural plasticity, rather than epistasis alone. Indeed, in contrast to this observation mutations which alter the chemistry and structure of the LBP in *h*ERs, are known to cause loss-of-function^25,31^. Our findings reveal an evolutionary exception to this rule and highlight differing levels of constraint within this key functional region that preserves ligand-dependent gene regulation by E2.

### Epistasis and degeneracy maintain homodimer interactions

The dimerization interface differs from the LBP and AF2 in that selective pressure arises from requirement to recognise self. Thus, to maintain interface compatibility co-evolution (epistasis) between opposing residues is likely required^36,37^. In ERα and ERβ homodimers, interface symmetry between subunits is maintained despite widespread sequence variation. Therefore, asymmetry of the *rf*ERγ dimerisation interface is unexpected. Structural analysis shows that this change stems from two opposing residues located above the hydrophobic patch, F497 and Y501, which are substituted for distinct amino acids between homologs (**Fig. 1g**). In the *rf*ERγ homodimer, Y501_a_ sterically displaces Y501_b_ into a constrained rotamer orientation establishing a hydrogen bonding network with the sidechains of D484_a_, the N^ε^ atom of R498_b_, T502_b_ and a water molecule. Concurrently, the Y501_a_ phenol also packs against F497_b_ and positions the phenyl ring for π-π stacking with F497_a_ (**Fig. 1g**). These alternate conformations mechanically leverage the two subunits apart, which would break a conserved hydrogen bond between D484 (H9) and Q498 (H10) maintained in ERα and ERβ homodimers (**Extended Data Fig. 3a**). However, in the *rf*ERγ homodimer, substitution for R498 rescues interaction with D484 due to the increased length of the sidechain and stronger ionic bond (**Fig. 1g**). We searched for further evidence of epistasis and found residues within the dimerisation interface that gain or lose interactions between homologs (**Extended Data Fig. 3b**). While structural plasticity does not directly result from these changes, epistasis between residues and substitution by ordered water networks (see *h*ERβ), act as compensatory mechanisms to variation. Moreover, previous work has shown that integrity of the hydrophobic patch is important for homodimerisation and confers strong selective pressure^15,39^. The sequence variation between homologs alters structure of the hydrophobic patch. However, substitutions are degenerate and maintain hydrophobicity while accommodating plasticity of interface structure. Together, these findings show that homodimerisation is preserved through a combination of degeneracy and epistasis, culminating in a varied spectrum of constraint within this key functional region.

### Allostery and protein folding are highly constrained

Allostery is essential for ligand-dependent regulation of ER LBDs; however, the architecture of the networks that transmit this information remains poorly understood. It is also unclear how allosteric mechanisms might constrain evolutionary variation and the implications for human diseases. Elucidating these networks is complicated by immense variability within a dynamic ensemble, let alone between five homologs with diverse sequences. To address this challenge, we integrated supervised-learning and comparative Markov modelling to resolve discrete conformational ensembles from the MD simulations (Methods; **Fig. 2a** and **Figs. S2-8**)^40,41^. We then applied a reductionist approach to identify ‘metastable’ residue interactions representing evolutionarily conserved network structures. The Markov models revealed that each homolog traverses a rugged conformational landscape at equilibrium (**Extended Data Fig. 4a**). Coarse graining the models by Perron-cluster cluster analysis revealed the most significant differences to be the conformational diversity of H6, H7, the base of H11 and the H12-loop, as well as mean-first passage timescales between macrostates (**Extended Data Fig. 4b**). These findings indicate that evolutionary changes in sequence can shift the conformational equilibrium of a structurally conserved domain. However, E2 and H12 remain stable indicating strong allosteric coupling (**Fig. S2**).

**Fig. 2.**
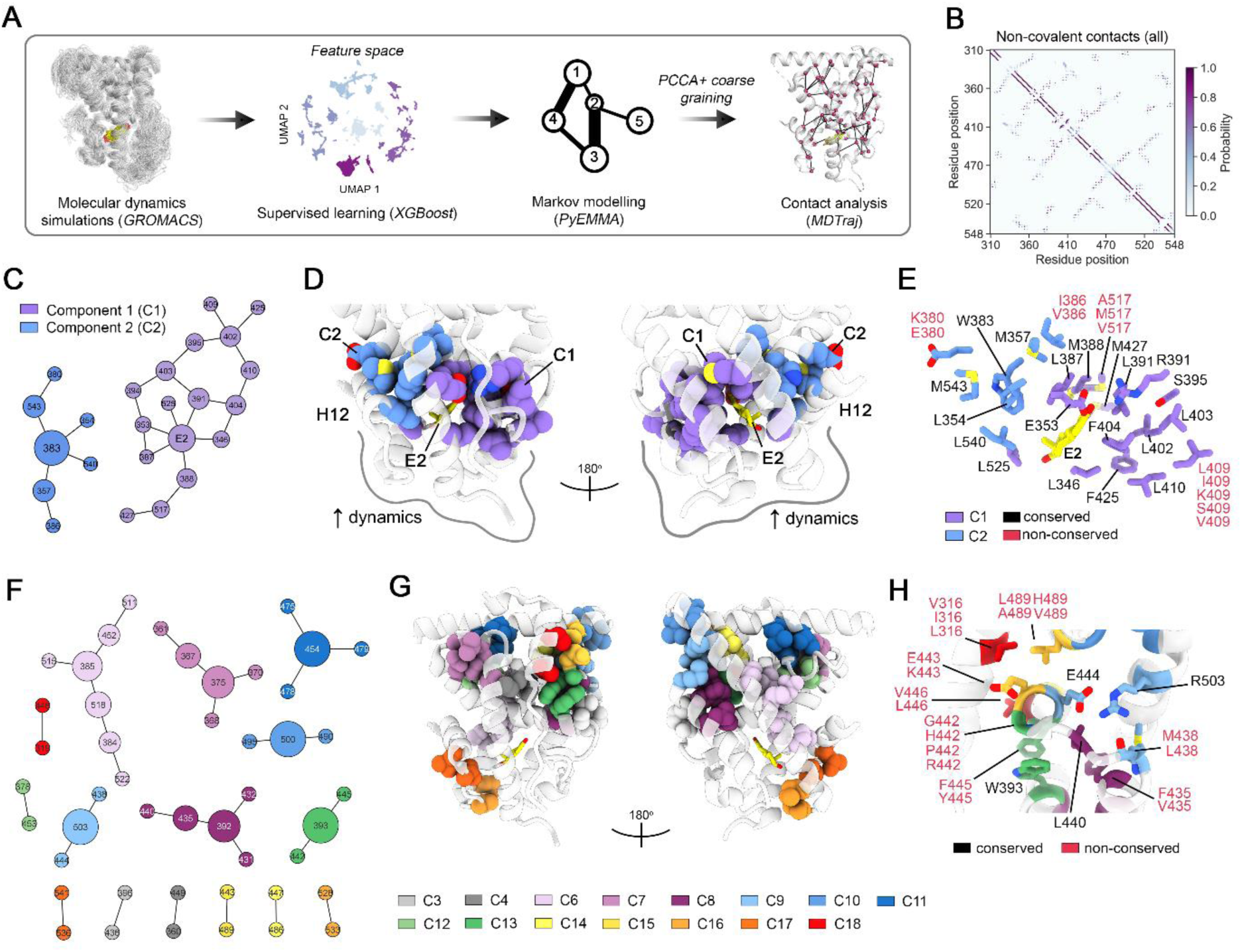
Identification of evolutionarily conserved network structures using an integrative computational approach. **a**, Schematic outlining the approach. See Materials and Methods for a detailed description and breakdown of each step. **b**, A contact matrix showing all non-covalent residue interactions between residues (all homologs) coloured by probability. **c**, Graph of the allosteric network comprised of components 1 and 2 (C1: purple; C2: blue). Node size is proportional to the betweenness centrality (or importance) of each node. **d**, Allosteric network mapped to the crystal structure of the wildtype *h*ERα LBD (PDB: 1GWR). Residue sidechains are shown as spheres and coloured by component. **e**, Structural analysis of the allosteric network. Conserved residues shared are labelled in black while non-conserved residues are labelled in red with varying amino acids annotated. **f**, Same as in **c** but for the components deemed important for supporting protein folding. **g**, The folding network mapped to structure. Residue sidechains are shown as spheres and coloured by component (see key). **h**) Perspective of the α-helical bundle and folding network components. Residue annotation follows convention of **d**.

We then calculated heavy-atom contacts between all residues, including E2 (≥ 90% occupancy and not within ± four positions along the protein backbone), from macrostates to identify network structures (**Fig. 2b**). These contacts were merged to produce a graphical network comprising multiple components that capture evolutionarily conserved, long-lived interactions over all ensembles. In this network, a component consists of a collection of nodes, where each node represents an individual residue position, and the edges represent non-covalent interactions between them. We mapped each component to the corresponding ER LBD structure and found that two components allosterically couple H12 to E2 and the LBP (**Fig. 2c**). The largest component, C1, forms a cluster of interactions between E2, H3, H5, the β-sheet, the centre of H7, and H11 (**Fig. 2d**). This architecture aligns with the stabilisation of *h*ERα and *h*ERβ LBDs upon E2 binding, as revealed by previous hydrogen-deuterium exchange coupled mass-spectrometry (HDX-MS) analyses^17,18,21^. The second component, C2, anchors H12 to H3, H4 and H5 through the formation of a hydrophobic cluster that converges around W383. C1 and C2 appear to work in tandem, forming an evolutionarily conserved architecture that facilitates both long- and short-range allosteric communication.

To assess the functional implications of this allosteric network in humans, we examined characterised mutations in *h*ER LBDs, focusing on those that would not directly perturb E2 binding to avoid biasing the analysis and masking of potential mechanisms. The analysis revealed that mutations disrupting the structure of the allosteric network cause LoF^42,43^. For instance, the F425S mutation likely decouples interactions with the β-sheet, while W383R and L540Q disrupt formation of the hydrophobic cluster. Sequence analysis of *h*ER and *rf*ER LBDs showed that key nodes in the allosteric network are conserved, while residues involved in mainchain interactions or solvent exposure are non-conserved (**Fig. 2e**). Definition of conserved allosteric network architecture provides a deeper understanding of the constraints of ligand-dependent regulation between vertebrates.

This network analysis also revealed constraints that govern folding of the ER LBD. We analysed other components identified by the contact analysis and found that each are small (on average ∼3 residues) and predominantly located within the α-helical bundle (**Fig. 2f** and **g**). For example, a small component comprised of three residue positions L/M438, E444 and R503, links H7 to H8 via H10 (**Fig. 2h**). A salt-bridge is formed between conserved residue positions E444 and R503, and the latter also hydrogen bonds with the carbonyl of L/M348. In the *h*ERα LBD, mutation of R503 to leucine disrupts the L/M348 carbonyl hydrogen bond and E444 salt-bridge causing LoF via destabilisation^42^. Similarly, mutation of other conserved residue positions within these components are destabilising in the *h*ERα LBD^44^. However, many of the non-conserved residues comprising each component are degenerate and maintain constrained interactions. Collectively, we propose that these components form the basis of an evolutionarily conserved scaffold that supports folding.

### Molecular constraints of the ER LBD are maintained between vertebrates

To investigate whether the molecular constraints identified in our analysis reflect natural sequence diversity, a comparative evolutionary analysis on 1,051 ERα, ERβ and ERγ LBD sequences from diverse vertebrates was performed. Ligand-independent protostome sequences were included as evolutionary and functional outgroups^45–47^. Evaluation of the variability within functional regions showed that the LBP is least varied (AUC = 0.930), followed by the AF2 (AUC = 0.893), the allosteric network (AUC = 0.861), the dimerisation interface (AUC = 0.707), and lastly the folding network (AUC = 0.685) (**Fig. 3a**). Indeed, residue positions encoding the LBP and AF2 are more conserved than non-functional regions per Jensen-Shannon divergence (JSD), with residues that specifically coordinate with conserved ligands, E2 and the coactivator *LxxLL* motif, being particularly more constrained (**Fig. 3b** and **c**). Structurally integral residues within the allosteric network are significantly more constrained compared to those that are degenerate and/or solvent exposed in our crystal structures. In contrast, residue positions encoding the dimerisation interface and protein folding network are non-significant. Structural analysis shows that residues cluster into pockets of varying conservation that correspond to changes observed within *h*ER and *rf*ER interfaces (**Fig. 3d** and **Extended Data Fig. 3b**). Network analysis indicates that the LBP, AF2 and allosteric network evolved neutrally around consensus genotypes with variation correlating to degenerate structural changes (**Fig. Extended Data Fig. 1b-d** and **5a-e**). Although, possible coevolution between AF2 residues and coactivators cannot be excluded. In contrast, the dimerisation interface is distributed into a sparse network of phylogenetically related communities, suggesting that, consistent with our structural findings from mammalian and teleost receptors, epistasis and degeneracy preserve homodimer assembly across vertebrates^34–37^. A limitation of our analysis is depth of species coverage in available genome data which could explain absent connectivity between some nodes. Alternatively, epistatic interactions might lead to complex mutational pathways that over time obscure connections between close phylogenetically related genotypes. Nonetheless, these findings show that ligand-dependent regulatory mechanisms and protein folding are conserved across vertebrates, revealing molecular constraints driving the structural evolution of a key transcriptional regulatory domain and adaption within functional regions to accommodate different selective pressures.

**Fig. 3.**
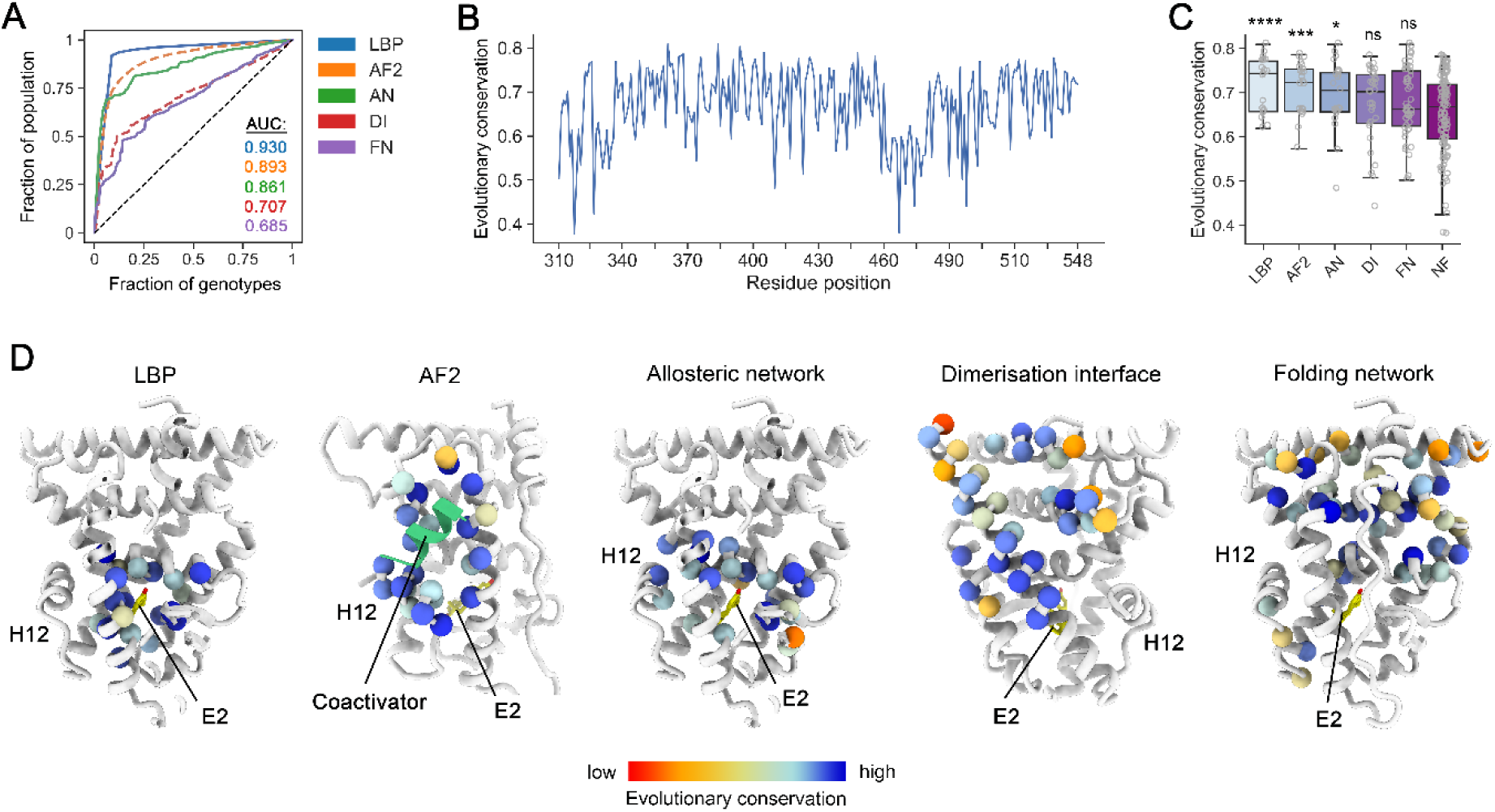
Evolutionary sequence analysis demonstrates that constraints are preserved across vertebrates. **a**, Evaluation of variability within functional regions of 1,051 ER LBD sequences (α, β and γ) from phylogenetically diverse taxa. Area under the curve (AUC) was used as a quantitative measure of constraint, calculated by plotting the fraction of unique genotypes in the alignment encoding functional regions (e.g., the dimerisation interface) against the fraction of the total sequence population. **b**, The evolutionary conservation of each residue position along the length of the LBD determined by the Jensen-Shannon divergence (JSD). **c**, Evolutionary conservation of each residue position encoding the ligand binding pocket (LBP), activation function-2 (AF2), allosteric network (AN), dimerisation interface (DI), folding network (FN) and non-functional residues (NF), shown as a box plot. The box plot shows the median and interquartile range (IQR); whiskers represent the distribution of the data as a function of the IQR. Mann-Whitney U-test was used to determine statistical significance compared to NF residues (LBP: *P* = 0.00095; AF2: *P* = 0.00303; AN: *P* = 0.03354; DI: *P* = 0.12424; FN: *P* = 0.10295). **d**, The evolutionary conservation of each residue position mapped to the *h*ERα LBD (1GWR; missing loops were modelled in ICM-Pro Molsoft for visualisation purposes) highlights the architecture of constraint within the ER LBD. Each residue position comprising regions are shown as spheres (Cα atom) and coloured according to the level of conservation (red = low; blue = high). Refer to **Extended Data Fig. 5** for the network analysis.

### The architecture of genetic variation in the human ER LBDs

Large scale genome sequencing has revealed widespread genetic variation within the human population^48,49^, yet much of the variation in human ERs and implications for disease remain uncharacterised. A clear picture of this landscape and the potential conflicts with constraints that preserve transcriptional regulatory mechanisms would be beneficial toward safe implementation of precision medicine strategies to treat diseases^50–52^. To investigate this, we analysed whole genome sequencing data for rare missense variants (<0.01% allele frequency) in *h*ERα and *h*ERβ LBDs from 807,162 individuals in gnomAD^48^, and compared them against the molecular constraints conserved across vertebrates. We identified 156 unique *h*ERα LBD variants at 108 residue positions, and 322 unique *h*ERβ LBD variants at 187 residue positions (**Fig. 4a,b** and **Extended Data Fig. 6a**). The variants span residue positions with diverse solvent accessibility, dynamics, and evolutionary conservation, yet trend toward structurally constrained positions (**Fig. 4c** to **e**). In the *h*ERα LBD, hypergeometric tests reveal that only the LBP and allosteric network are significantly depleted of variants (**Fig. 4f,g** and **Fig. S9**), demonstrating that hormone binding and allosteric regulation are functionally constrained and under strong selective pressure. In contrast, *h*ERβ displayed no significant depletion in any functional region indicating widespread tolerance to variation. However, structural analysis of both receptors revealed widespread disruption of constraints and the existence of known LoF variants, such as W383R and R394H; located within the allosteric network (**Fig. 4g,h**). Other examples are mutation of R503 in the protein folding network to the variants R503Q and R503W in *h*ERα, and R503C and R503H in *h*ERβ; each variant would disrupt the constrained ionic/hydrogen bonding network linking H7, H8 and H10, causing destabilisation and LoF^42^. However, in the LBP, the *h*ERα variant, M421V, is likely to be benign given the relaxed constraint at this position and is a degenerate change. We speculate that differences in variant burden and architecture between receptors arise from their evolutionarily conserved but diverse physiological roles, particularly for *h*ERα in reproductive biology. However, the pervasive disruption of constraints would suggest that many individuals harbouring variants could have impaired function in both receptors^15,23,25,42,43^.

**Fig. 4.**
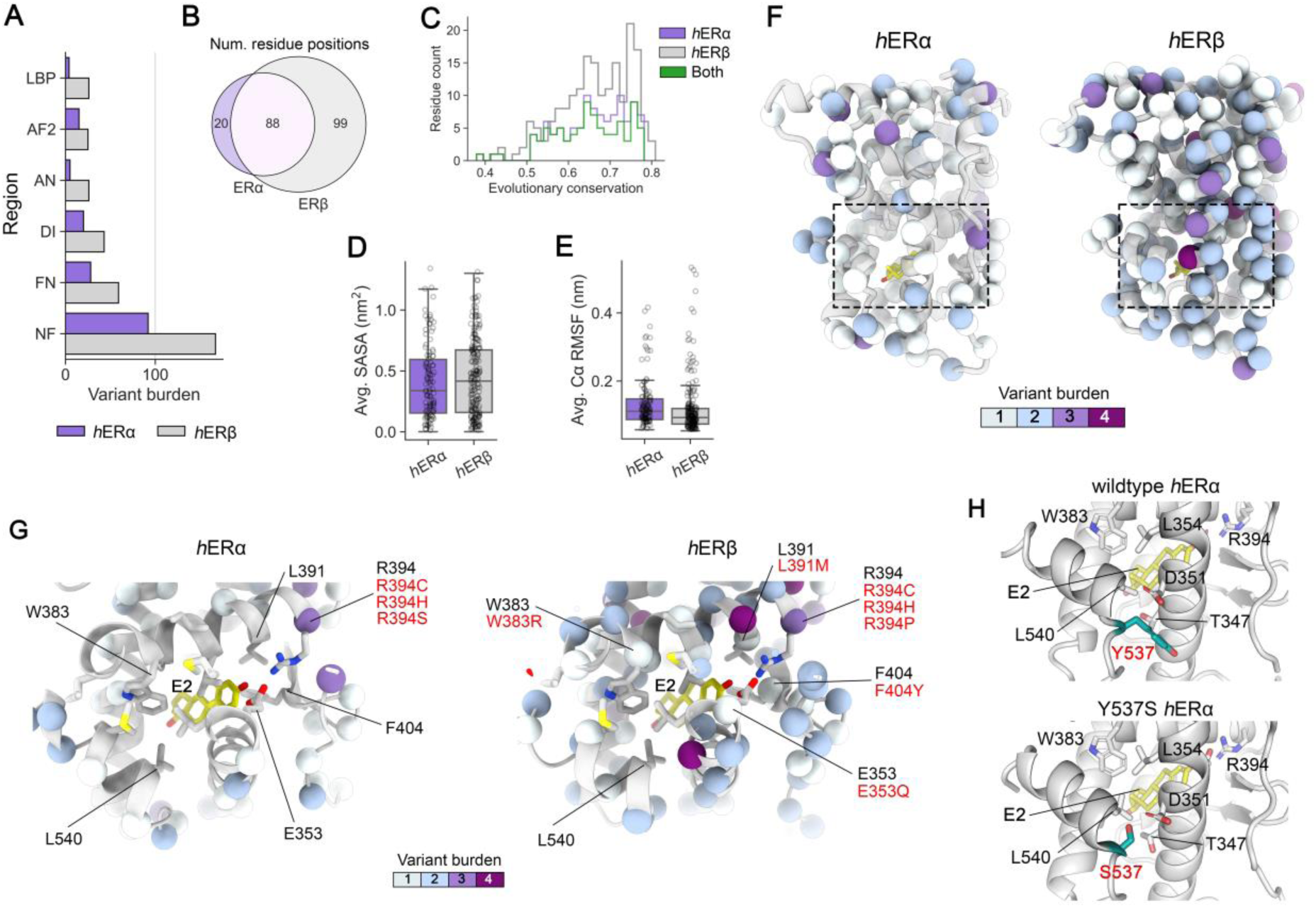
Human genome sequencing data (gnomAD) reveals pervasive variation in hERα and hERβ LBDs and widespread disruption of constraints. **a**) The burden, or total number, of unique variants within each functional region shown as a grouped barplot (LBP: ligand binding pocket; AF2: activation function-2; AN: allosteric network; DI: dimerisation interface; FN: folding network; NF: non-functional residues). Note that a residue position may harbour more than one unique variant. **b**, Venn diagram showing the distribution of shared residue positions harbouring variants between *h*ERα and *h*ERβ LBDs. **c**, Histogram showing the distribution of the evolutionary conservation (Jensen-Shannon divergence) of each variant residue position for both *h*ERα and *h*ERβ, as well as residue positions that share variant occurrence. **d**, Average solvent accessible surface area (SASA) of residue positions harbouring variants calculated from the homodimeric crystal structures of the *h*ERα (PDB: 1GWR) and *h*ERβ (PDB: 3OLL) LBD. Missing residues in loops were modelled with ICM-Pro Molsoft. **e**, Average Cα-atom root-mean squared fluctuations (RMSF) of residue positions harbouring variants calculated from the replicate MD simulations. **f**, Molecular architecture of residue positions harbouring variants mapped to *h*ERα (PDB: 1GWR) and *h*ERβ (PDB: 3OLL) LBDs. Residues are shown as spheres (Cα atoms) and coloured according to variant burden. The black box highlights the differences between variant burden in ligand binding pocket and of allosteric network residues between *h*ERα and *h*ERβ LBDs. Whiskers in **d** and **e** represent median and interquartile range (IQR). **g**, A structural perspective of the ligand binding pocket and allosteric network residues of the *h*ERα LBD (left) and *h*ERβ LBD (right). Important residue sidechains are shown as sticks. Residue positions with variants are identified by a sphere which is coloured to the total burden of unique variants. Wildtype residues are labelled in black, while identified variants are labelled in red to highlight changes. **h**, Structural comparison between wildtype *h*ERα (PDB: 1GWR) and Y537S (gain-of-function; PDB: 3UUD). Adjacent allosteric network residues are shown as sticks and the residue at position 537 is coloured teal

ER+ breast cancers are primarily driven by mutations, such as Y537S and D538G, that constitutively activate the *h*ERα LBD, promoting uncontrolled cellular proliferation and tumour development^20,22^. Why these variants occur and at their specific location within the AF2 is unclear. Constitutively activating variants were not identified in the genome sequencing data, likely due to their somatic origins and long-term negative outcomes. Structurally, these variants cluster near, but outside, the allosteric network, stabilising H12 in the active conformation and lowering the energy penalty to activation (**Fig. 4h**). Constitutive activation arises through positive epistasis with residues on H3 and relaxation of structure which improves packing^21,22,53^. Our findings provide new insight, demonstrating that the constraints which deplete variants from the allosteric network select for mutations that enhance allosteric regulation, ultimately driving constitutive activity and disease. These findings highlight the context specific effects of mutations and how nature can strike a fine balance between optimal protein fitness in normal conditions and deleterious gain-of-function outcomes.

In summary, this work provides new insight into the mechanisms that maintain functions of a key transcriptional regulatory domain and highlight the power of comparative analysis to address fundamental biological questions. These findings further demonstrate that the origins of natural genetic variation in humans, and ER- related diseases, are directly linked to the molecular constraints that protect gene regulation during vertebrate evolution. We anticipate that similar analysis of other transcription factors will enhance understanding of gene regulation across nature and disease origins.

## Materials and Methods

### Materials and Methods

#### Expression and purification of recombinant proteins

In this study, wildtype ERα, ERβ and ERγ LBDs from *Melanotaenia fluviatilis* (Murray River rainbowfish; *rf*ERs)^27^ and ERα and ERβ LBDs from *Homo sapiens* (humans; *h*ERs)^4,5^ were recombinantly expressed in *Escherichia coli* and purified by fast protein liquid chromatography (FPLC) using an NGC Medium-Pressure Liquid Chromatography system (*Bio-rad*).

##### Expression of recombinant protein

Codon-optimised *rf*ERα, *rf*ERβ and *rf*ERγ LBD primary sequences were cloned into pET-21b (*Twist Bioscience*); *h*ERα and *h*ERβ LBD primary sequences were cloned into pET-11a vector (*Genscript*). Each construct contained a non-removable N-terminal hexa-histidine tag. Constructs were transformed into *Escherichia coli* BL21(DE3) for recombinant expression. The following day, an isolated colony was used to inoculate 100 mL LB medium supplemented with 100 μg/mL ampicillin, which was then incubated overnight at 37°C. Next, seed culture was diluted 1:200 into 1L flasks containing LB medium supplemented with 100 μg/mL ampicillin and cultured at 37°C at 180 rpm until OD_600_ = 0.5-0.8 upon which 500 μM IPTG were added to induce expression of recombinant protein. Where necessary, cultures were treated with 5-10μM 17β-estradiol (E2) for co-fermentation. For the fluorescence polarisation (FP) assays, the *rf*ERα LBD and *h*ERβ were purified apo, in the absence of ligand. Recombinant protein was expressed for 20-24hrs at 16°C before harvesting at 4,000-5,000xg for 20 minutes.

##### Purification of recombinant protein for crystallography

All steps herein were performed at 4°C or on ice. Cell pellets were resuspended in 25 mL buffer A (20 mM Tris-HCL pH 7.5, 500 mM NaCl, 10 mM imidazole, 10% glycerol, 5 mM β-mercaptoethanol (BME), 0.1% Tween-20), pooled and gently stirred in a beaker. To the cell suspension hen egg-white lysozyme crystals were added. The lysis reaction was incubated for 30 minutes to 1 hour (depending on volume) and then lysed by mechanical disruption. The lysate was then centrifuged at 40,000xg for 1hr to fractionate and clarify the sample. The supernatant was then filtered with a 0.45 μm cut-off syringe filter. Filtered lysate was loaded over an equilibrated 5 ml HisTrap HP column (*Cytiva*), washed with 50 ml buffer A (minus tween-20) and eluted in buffer B (20 mM Tris-HCL pH 7.5, 500 mM NaCl, 250 mM imidazole, 10% glycerol, 5 mM BME) over a gradient. Fractions containing enriched protein by SDS-PAGE analysis were diluted 10-fold immediately into buffer C (20 mM Tris-HCL pH 7.5, 10% glycerol, 5 mM BME) and further purified by cation exchange chromatography using a 5 ml HiTrap Q HP column (*Cytiva*) and eluted over a linear percentage gradient of increasing buffer D (20 mM Tris-HCL pH 7.5, 500 mM NaCl, 10% glycerol, 5 mM BME). Proteins typically eluted in the 150-200 mM NaCl range. For X-ray crystallography, fractions were pooled and concentrated to less than 10 mL then further polished by size exclusion chromatography using a HiPrep 26/60 S-200 HR column (*Cytiva*) and eluted into buffer E (20 mM Tris-HCL pH 7.5, 200 mM NaCl, 10% glycerol, 5 mM BME). The elute was concentrated to ∼10-15 mg/mL and flash-cooled in liquid nitrogen for storage at −80°C.

##### Purification of recombinant protein for fluorescence polarisation coactivator binding assays

All steps herein were performed at 4°C or on ice. As above, cell pellets were resuspended in 25 mL buffer A and lysed by mechanical disruption, then clarified by centrifugation at 40,000xg for 30 minutes. The clarified lysate was loaded over a 5 ml HisTrap HP column (*Cytiva*) equilibrated with buffer E (20 mM Tris-HCl pH 8.0, 500 mM NaCl, 10 mM imidazole, 2 mM BME, 10% glycerol), washed with 50 ml buffer E, and eluted in two steps of 25 ml 65% and 100% buffer F. (20 mM Tris-HCl pH 8.0, 500 mM NaCl, 250 mM imidazole, 2 mM BME, 10% glycerol). Fractions containing enriched protein by SDS-PAGE analysis were diluted 8-fold immediately into buffer G (20 mM Tris-HCl pH 8.0, 10% glycerol, 5 mM DTT) and loaded over a 5 ml HiTrap Q HP column (*Cytiva*) equilibrated in buffer F (20 mM Tris-HCl pH 8.0, 100 mM NaCl, 10% glycerol, 5 mM DTT). Following sample loading the column was washed with 25 ml buffer D and eluted over a linear gradient of increasing elution buffer H (20 mM Tris-HCl pH 8.0, 500 mM NaCl, 10% glycerol, 5 mM DTT). Proteins typically eluted in the 150-200 mM NaCl range. The fractions containing pure protein were concentrated to >10mg/mL and flash-cooled in liquid nitrogen for storage at −80°C. Note: in both purification strategies *h*ERβ and *rf*ERα LBD were purified apo but *h*ERα, *rf*ERβ and *rf*ERγ LBDs were insoluble without co-fermentation and co-purification with E2. Protein concentrations for both methods were measured with a ThermoFisher NanoDrop using molecular weights and theoretical extinction coefficients.

#### Fluorescence polarisation (FP) coactivator binding assays

To perform FP coactivator binding assays a previously described peptide probe was used (D22; 5-FAM-LPYEGSLLLKLLRAPVEEV-COOH; 98% purity), and was purchased from *GenScript* (Piscataway, NJ, USA)^54,55^. For FP assays a 200 µM E2 stock was prepared in 100% DMSO. The coactivator probe was prepared as a 10 mM stock in MQ H_2_O and adjusted to ∼ pH 7 with dropwise addition of 2M NaOH, with working stocks subsequently prepared at a final concentration of 1 µM in 10 mM Tris-HCl pH 7.5 and stored at −20°C.

All fluorescence polarization experiments were performed using the same general procedure at 22 °C with samples (160µl final volume) containing FP Buffer (20 mM Tris-HCl pH 8.0, 100 mM NaCl, 0.05% Tween 20, 0.1 mM TCEP, 10 µM estradiol, 5% DMSO and 10 nM D22 probe), with the concentrations of the ER LBD variant varied. Once prepared, samples were incubated at 22 °C for 30 min. Following incubation, 150 µl of each sample was transferred to a single well of a non-binding flat-bottom black 96 well plate (Greiner Ref. 655900). Fluorescence polarization was measured using a Pherastar FX microplate reader equipped with the FP 485 520 520 module using default settings. mP values were calculated using Equation 1 following subtraction of raw parallel and perpendicular emission values for a corresponding sample with the coactivator probe omitted.

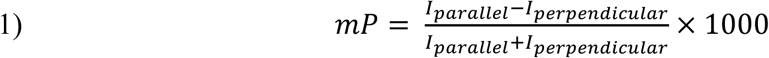

To study the interaction of the coactivator class probe with the ER LBD in the presence and absence of E2, samples contained FP Buffer and 8 concentrations of ER LBD homolog ranging between 0.3 and 5× the estimated dissociation constant (*K*_d_). The *K*_d_ was estimated by non-linear regression in GraphPad Prism using Equation 2, where *mP_free_* and *mP_bound_* are the fluorescence polarization for free probe and saturated receptor respectively, [*L*] is the total concentration of coactivator class probe, [*R*] is the total concentration of ER LBD homolog and *K_d_* is the dissociation constant.

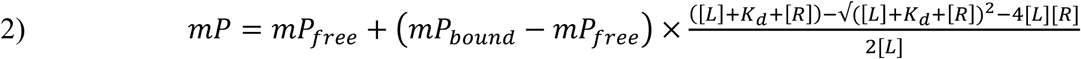

#### X- ray protein crystallography and modelling

Sparse matrix crystal screening was performed to identify initial lead crystallisation conditions. Sparse matrix crystal screens were obtained from *Hampton Research* and *Molecular Dimensions*. Each sparse matrix crystal screen was established in 96-well sitting drop trays (*Hampton* Research) with 80 µL reservoir solution and 1:1 ratio of protein:well-solution at 16°C. Manual optimisation was performed using the hanging-drop vapor diffusion method. The *rfER*α LBD (at 10 mg/mL) was co-crystallised with E2 (incubated with 1:1.5 molar ratio) and *h*SRC2-2 coactivator peptide (5-KHKILHRLLQDSS-OOH; incubated with 1:2 molar ratio) in 9-10% Tacsimate pH 7.0 and 22% PEG3000 at 16°C; crystal morphologies were cubic and grew to be 75-100 µm in diameter. The *rfER*γ LBD (at 10 mg/mL) was co-crystallised with E2 (1:1.5 molar ratio) in 0.2 M Magnesium formate dihydrate and 20% PEG3350 at pH 7.0 at 16°C following extensive optimisation; morphology of the single diffracting crystal was hexagonal/rod shaped and approximately 100-150 µm in length. Optimisation was performed by hanging-drop vapour diffusion with a reservoir volume of 500 µL. Unfortunately, only very weakly diffracting crystals could be obtained of the *rfER*β LBD even with extensive optimisation of crystallisation conditions including protein concentration, ligand to protein ratio, temperature, drop size, micro-seeding, and cross-seeding. In lieu, an energy minimised homology model was built with *ICM-Pro* using the *h*ERβ LBD as a structural template ^56^. Models of missense variants were generated with *ICM-Pro* followed by local energy minimisation and refinement.

#### Data collection, refinement, and analyses of crystallographic data

*Crystallographic data collection and refinement.* Protein crystals were cryo-protected in NVH oil and flash-cooled by plunging into liquid nitrogen prior to X-ray diffraction. Data collection was performed at the Australian Synchrotron on the MX1 and MX2 beamlines^57,58^. Diffraction data were collected over 360° at 0.1° oscillation. The *rf*ERα LBD co-crystal was solved at 2.75 Å in space group I4_1_32 and the *rf*ERγ LBD co-crystal was solved at 1.95 Å in space group P6_3_ (**Supplementary Table 1**). Crystallographic diffraction data were indexed, merged, and scaled in XDS and Aimless^59,60^. Molecular replacement (MR), refinement and model building were performed using *Phaser*, *PHENIX*, and *Coot*^61–63^. During MR, the crystal structures of human ERα and ERβ LBDs were used as search models for the *rf*ERα and *rf*ERγ LBDs, respectively. *Chainsaw* was used to prepare the search models ^64^. Additional analyses, such as root-mean squared deviations (RMSDs) between receptors and residue-residue distances were calculated using *Coot*^61^ and *PyMOL* (*v2.0 Schrödinger, LLC*). Solvent accessible surface area (SASA) was calculated using *MDTraj* and *NumPy* libraries^65,66^. Network topologies were visualised using *Cytoscape*^67^.

#### All-atom molecular dynamics (MD) simulations

All-atom MD simulations were performed for *h*ERα, *h*ERβ, *rf*ERα, *rf*ERβ and *rf*ERγ LBDs bound to E2 in explicit solvent at 310K for comparison. MD simulations were performed using GROMACS with the Charmm36 forcefield^68–71^. Ligand topologies were generated using the SwissParam server^72^. Each receptor was centred in the middle of a dodecahedral box no less than 1.0 nm from the periodic edge boundary and solvated with the TIP3P water model. When required, sodium and chloride ions were added to the system to neutralise net charge. The system was energy-minimised using the steepest-descent algorithm until *F_max_* < 1000 and then equilibrated by sequential 1 ns restrained NVT and NPT simulations using the Lincs constraint algorithm, Particle Mesh Ewald electrostatics, Berendsen thermostat coupling and Parrinello-Rahman pressure coupling (NPT only). Next, restraints were released, and 1 μs production simulations were performed in triplicate using a 2-femtosecond leap-frog indicator. Frames were recorded every 100ps for storage efficiency. All MD analyses and graphical visualisation were performed in *Python* using the *MDTraj, NumPy, Matplotlib* and *Seaborn* libraries^65,66,73,74^.

#### Markov State modelling

##### Feature selection

The dimensionality of a protein is immense, let alone over an ensemble and between homologs with different sequence. To identify features that capture structural and dynamical changes between homologs, pairwise distances were measured between Cα atoms for each frame of the concatenated trajectories. Excluded from the calculations were the disordered H8-H9 loop; the residues ∓ three at the N- and C-termini; and residue 328 (a gap in *h*ERβ). While dynamics of the H1-H3 loop are quite variable, hydrogen-deuterium exchange mass-spectrometry (HDX-MS) shows that the loop is stabilised upon ligand binding for the *h*ERα and *h*ERβ LBDs, and therefore it was not omitted^17,75^.

Next, we implemented a supervised machine learning approach to select features. First, the dataset was filtered to remove features with low variance (≤ 0.1) and then each feature was Z-score normalised. Initially, we trained eight different classification algorithms with 5-fold cross validation on the dataset (70:30 train-test split) with default hyperparameters: Gaussian Naïve Bayes (GNB), K-Nearest Neighbours (kNN), Linear Support Vector Machine (Linear SVC), Logistic Regression (LR; *max_iter* = 10,000), Linear Discriminant Analysis (LDA), Support Vector Machine (SVM), and RandomForest (RF) implemented using *Scikit-learn* (*sklearn*) ^76^ and Extreme Gradient Boosting (XGBoost^77^) decision tree classification algorithm in *Python*. Each model performed well (data not shown); however, because XGBoost is more resilient towards overfitting and has built-in methods to rank feature importance, we selected this algorithm to train on the dataset. To tune hyperparameters, RandomizedSearchCV was implemented using *sklearn.model_selection* with 5-fold cross-validation and ‘precision_weighted’ was used as the scoring metric and early-stopping was used to prevent overfitting. Final model parameters were: *subsample* = 0.5, *n_estimators* = 200, *max_depth* = 5, *learning_rate* = 0.01, *colsample_bytree* = 1. The model was evaluated on the test set (30% of the data which was randomly selected) by calculating the area under the receiver-operator characteristic curve (micro-averaged ROC-AUC = 1.0) and the area under the precision-recall curve (micro-averaged PRC-AUC = 1.0). Important features were identified directly from the hyperparameter optimised model using the *model.feature_importances_* function implemented in the *XGBoost* library^77^. In total, twenty features that capture long and short-range interactions were selected based on their ranked importance. To visualise the distribution of feature space, the top Z-score normalised features were discretised to two-dimensions using Uniform Manifold Approximation and Project (UMAP). All working described above was performed using the *MDTraj, NumPy, scikit-learn*, *UMAP* and *XGBoost* libraries^65,66,76–78^.

##### Markov State Model construction and validation

Markov state models (MSMs) for each homolog were built using *PyEMMA* software^79^. First, feature space was discretised to two-dimensions using time-structured independent component analysis (TICA) of the Z-score normalised features with a lag time *π* = 100 ps (1 step). To identify the appropriate lag time for each homolog a broad range was sampled, but we observed no discernible difference in discretisation as *π* was increased. Therefore, 100 ps was selected to resolve faster processes. The discretised trajectories were then clustered into microstates using K-means algorithm implemented in *PyEMMA*. To identify the optimal number of clusters, five unvalidated MSMs were estimated over an increasing range of *k* = 5 to 3000 and the VAMP-2 score was used as heuristic with 90% confidence interval. The optimal number of clusters was determined by the plateau of the curve. Bayesian MSMs were constructed for each homolog the number of macrostates were determined based on the convergence of implied timescales and statistically validated using the Chapman-Kolmogorov test. Macrostate assignments were coarse-grained using perron-cluster cluster analysis (PCCA) and mean-first passage times (MFPTs) were estimated using transition-path theory (TPT) implemented in *PyEMMA*^79^.

##### Contact enrichment analysis

To identify evolutionarily conserved networks, pairwise contact probabilities for the top fifty microstate assignments of each macrostate were calculated using *MDTraj*^65^ with a heavy-atom distance cut-off of ≤ 0.42 nm (4.2 Å) and not within four residues along the protein chain. This analysis was repeated for each homolog and then merged; a ‘metastable’ contact between two residues was determined if the occupancy of the interaction or probability was ≥ 0.9 or 90% occupancy. Graphical networks were constructed using *NetworkX* and visualised with *CytoScape*^67,80^.

#### Evolutionary sequence analysis

##### Sequence profile mining, maximum likelihood phylogenetics and evolutionary conservation

Homologous sequences were obtained from the UniRef-100 database using Hidden Markov Model (HMM) profile mining with *HMMER* v3.4 (http://hmmer.org/) and from the NCBI database using *BLAST*^81,82^. To build the HMM profile, experimentally validated primary sequences were first obtained from UniProt (https://www.uniprot.org/), iteratively aligned using *MAFFT v7.0* ^83^ and the positions manually refined according to human ERα and ERβ LBD structures (PDB IDs: 3UUD and 3OLL, respectively). The structure-guided alignment was used to build an HMM profile with *hmmbuild* (default parameters). Next, the UniRef-100 database was mined using the HMM profile with *hmmsearch* and merged/aligned to the structure-guided alignment with *hmmalign*. After filtering to remove redundant sequences and further refinement to retain sequences with less than 10% gaps the final alignment comprised 1,051 primary sequences of length 239 positions. Maximum-likelihood phylogenetic reconstruction was performed using *IQ-TREE2* with 10,000 ultrafast bootstrap replicates and hill-climbing nearest neighbour interchange optimization^84,85^. To identify the substitution model, JTT+R7, we used *ModelFinder* ^86^. The final ML-tree was rooted at the branching between deuterostomes and protostomes and analysed using *iTOL* (https://itol.embl.de/). To determine the evolutionary conservation of each residue position Jensen-Shannon divergence (JSD) was calculated in *SciPy* ^87^.

##### Graph network analysis of sequence space

Graphical networks were created for each functional region (e.g., the ligand binding pocket) to visualise the distribution of sequence space. New alignments were created from the primary alignment constituting positions for each functional region and then filtered to only include non-redundant sequences. Pairwise genetic distances between each sequence were calculated as the Hamming distance:

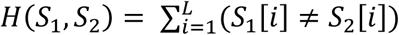

where *H*(*S*_1_, *S*_2_) is a summation of the number of positions between two sequences, *S*_1_ and *S*_2_, that differ in amino acid identity. Next, undirected graphs were constructed where two sequences (nodes) *i* and *j* are connected by an edge in the adjacency matrix *A*[*i*][*j*] if the Hamming distance between them is equal to 1 i.e., a single residue substitution. Attributes, such as sequence and count in the non-redundant alignment (population), were added to each node. To determine the shortest mutational path lengths between sequences, a connected graph was constructed where *A*[*i*][*j*] is the weight of the edge according to the Hamming distance between nodes *i* and *j*. Dijkstra’s algorithm was implemented to find shortest mutational path lengths between pairs of sequences using genetic distances as edge weights^88^.

##### Sequence constraint

To determine constraint for each functional region, area under the curve (AUC) was calculated: a high AUC indicates strong constraint throughout sequence space whereas a low AUC indicates less constraint. Here, the curve is calculated by plotting the cumulative fraction of the population (of the sequence alignment) against the number of unique genotypes indexed by decreasing frequency. AUCs were calculated using *Scikit-learn*.

#### Analysis of human genetic variation

To identify rare genetic variation of ER genes among humans (≤ 0.1% allele frequency), missense variation data was obtained from the Genome Aggregation Database (gnomAD *v4.1.0*, https://gnomad.broadinstitute.org/). Curation of variants was performed manually. Premature terminations were not included in the analysis, and hypergeometric tests were performed using *SciPy*. Visualisation of variant data was performed in *Python* using *Seaborn* and *Matplotlib* libraries^73,74^.

## Extended Data Figures

**Extended Data Figure 1.**
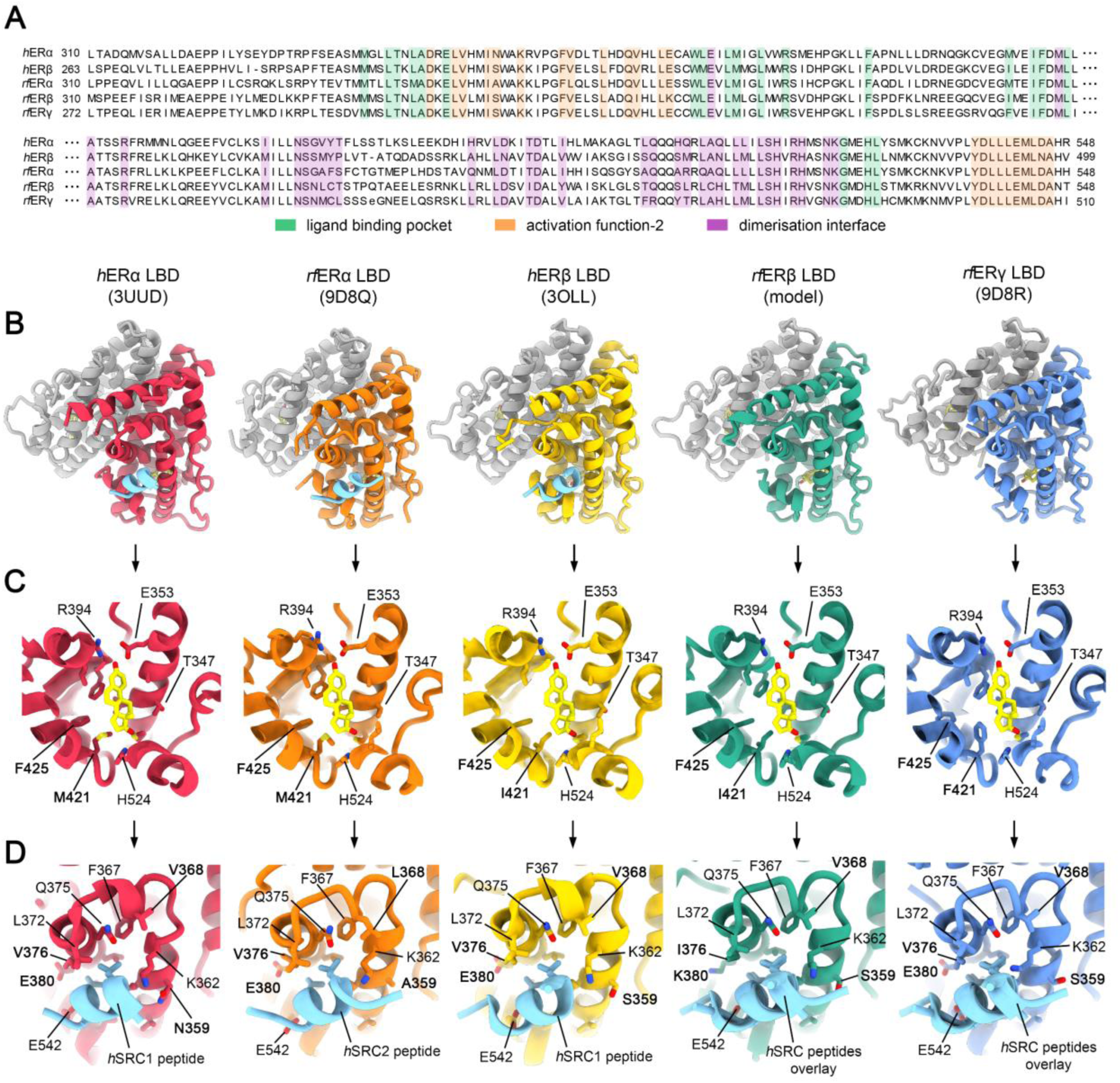
Structural comparisons between hER and rfER LBDs. a, An alignment comparing the primary amino acid sequences of each homolog. The numbering at sequence ends are the start and finish residue positions of each LBD; however, herein *h*ERα LBD numbering is used for consistency. Residue positions encoding the ligand binding pocket (LBP) are shown in green; the activation function-2 interface (AF2) in orange; and the dimerisation interface shown in purple. b, X-ray crystal structures of the *h*ERα LBD bound to estradiol (E2) and *h*SRC1 coactivator peptide (PDB: 3UUD); the *rf*ERα LBD bound to E2 and *h*SRC2 coactivator peptide (PDB: 9D8Q); the *h*ERβ LBD (PDB: 3OLL) bound to E2 and *h*SRC1 coactivator peptide; a computational homology model of the *rf*ERβ LBD bound to E2; and X-ray crystal structure of the *rf*ERγ LBD (PDB: 9D8R) bound to E2. Each structure is shown as the homodimer, with the second subunit coloured in grey. c, Close-up perspective of the LBP showing the conservation of E2 binding mode across homologs. d, The binding mode of coactivator peptides solved with *h*ERα, *rf*ERα and *h*ERβ, and all peptides modelled onto the AF2 of *rf*ERβ and *rf*ERγ LBDs, revealing conservation of structure and *LxxLL* motif recognition.

**Extended Data Figure 2.**
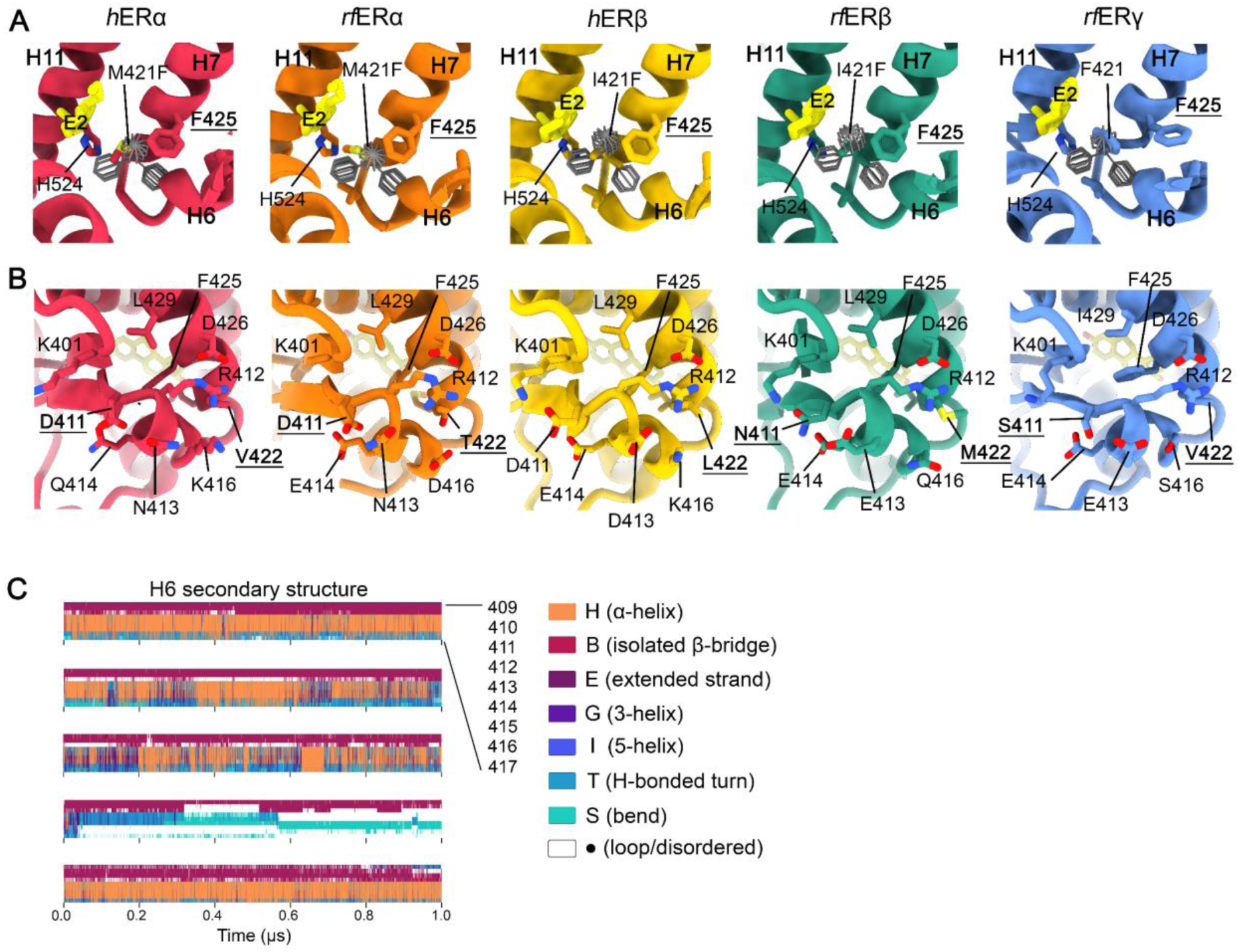
Structural analysis of the F421 substitution and surrounding environment. a, Modelling of all possible rotamer orientations of the F421 substitution in ERα and ERβ backgrounds with the Dunbrack rotamer library^89^ shows clashing of the phenyl sidechain with E2 and neighbouring structure, compared to the wildtype amino acid. b, Comparison of the structure and sequence of H6 between homologs. c, Secondary structure analysis of H6 (residues 409 to 417) calculated from a 1 µs all-atom molecular dynamics (MD) simulation using the DSSP algorithm in MDTraj ^65,90^.

**Extended Data Figure 3.**
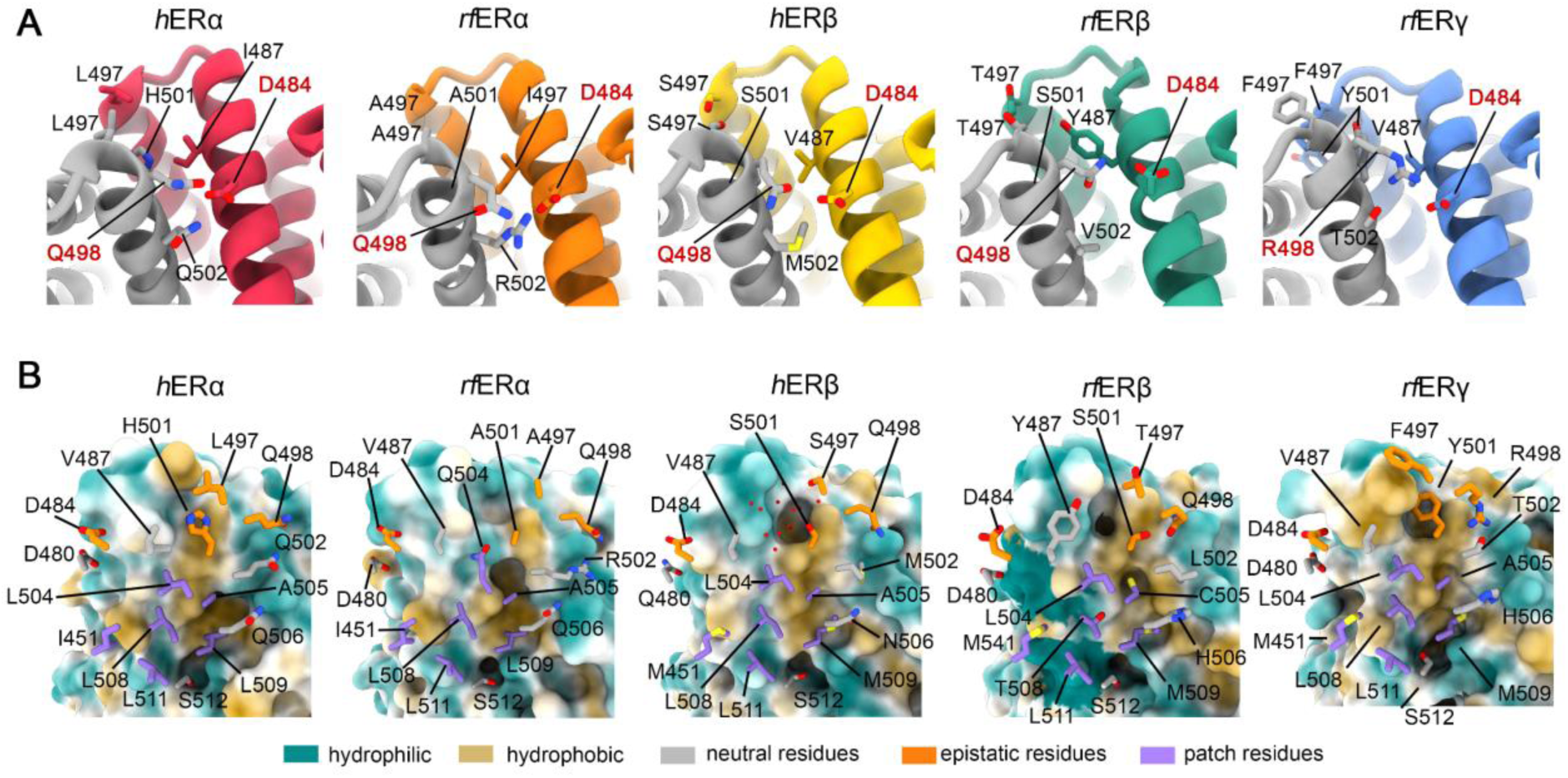
Comparison of ER LBD dimerisation interfaces. a, A close-up perspective of the top of the dimerisation interface highlighting amino acid differences and the constrained interaction between residue positions D484 and Q/R498. b, Structural and evolutionary comparison of the dimerisation interface reveals the existence of degeneracy and epistasis to maintain homodimer assembly. A surface representation of one subunit is shown and coloured by hydrophobicity/hydrophilicity; a subset of opposing important dimerisation interface residues are shown as sticks and coloured according to attribute: neutral residues (grey), epistatic residues (orange) and hydrophobic patch residues (purple).

**Extended Data Figure 4.**
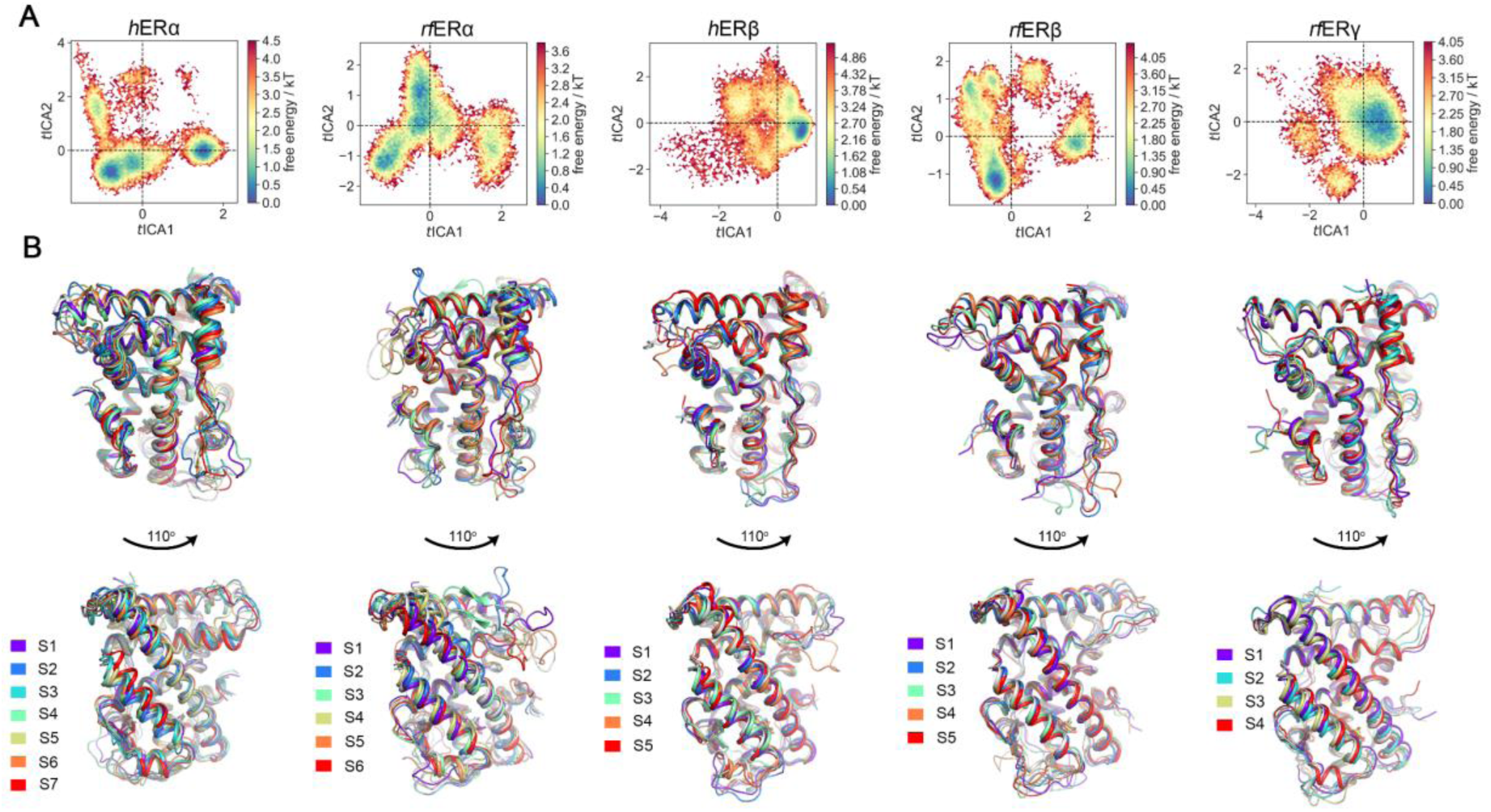
Evolutionary variation reshapes the conformational landscape between homologs. a, Free-energy surface plots for each homolog calculated during Markov State model construction in PyEMMA ^79^, mapped onto the first two components of the time-structured independent component analysis (TICA) plot. b, Representative cluster centroids from microstate assignments of each macrostate obtained from coarse-graining MSMs, coloured by membership. Front and rear perspectives of the LBD are shown.

**Extended Data Figure 5.**
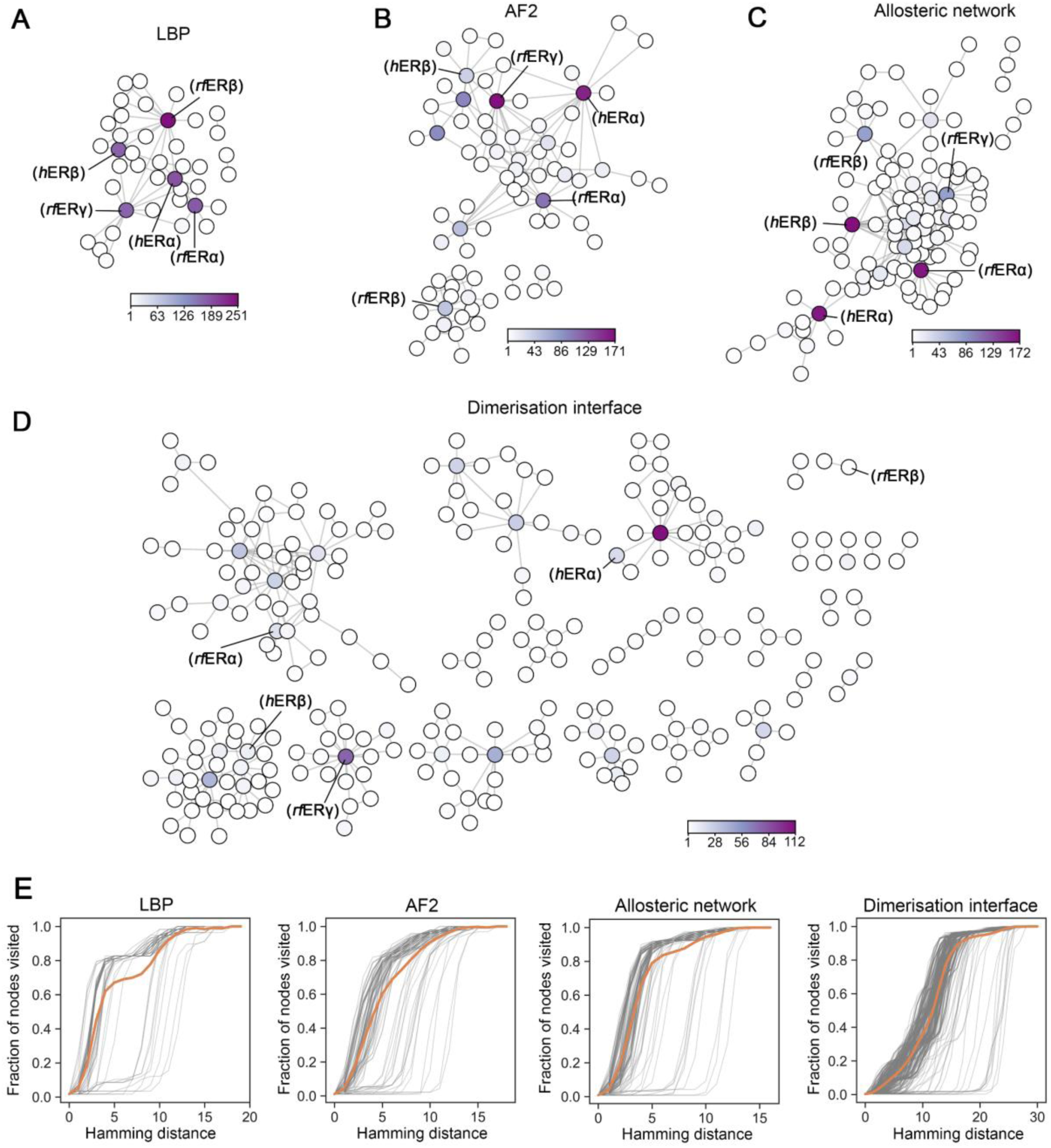
Network analysis of the sequence space encoding each functional region. a-d, Sequence space networks of the ligand binding pocket (LBP), activation function-2 interface (AF2), allosteric network and dimerisation interface. A genotype is depicted as a node and are connected by an edge to another node if the Hamming distance between them is equal to 1, or 1 residue substitution (i.e., the Hamming distance between AGFST and AFFST is 1). The colour of each node corresponds to the number of times each genotype is observed in the alignment of 1,051 ER LBD sequences. For visual clarity only connected nodes are shown. e, The number of mutational steps that each genotype (a line) must take to visit all other genotypes in the network, given as the Hamming distance, or shortest-mutational paths. The solid orange line represents the mean trajectory.

## Acknowledgments

This research was undertaken in part using the MX1 and MX2 beamline at the Australian Synchrotron, part of ANSTO, and made use of the Australian Cancer Research Foundation (ACRF) detector. Computational resources were provided by the University of Adelaide Phoenix High-Performance Computing (HPC) facility. We thank P. Bain and A. Kumar for discussion regarding *M. fluviatilis* sequences; T. Ritchie for assistance with initial optimisation of the *rf*ERα LBD purification; F. Voisin for Phoenix HPC support, and members of the Bruning Laboratory for constructive discussion. The authors acknowledge the use of Chat-GPT for identification of grammatical errors. The authors also thank D. Peet, M. Whitelaw, M. Hutchinson and K. Dholakia for comments on the manuscript.

## Funding

D.P.M. was supported by an Australian Government Research Training Program (RTP) scholarship. This work is supported by an Australian Research Council Discovery Project (DP230100609) and the University of Adelaide Biochemistry Trust.

## Author contributions

Conceptualization: D.P.M. and J.B.B. Funding acquisition: J.B.B. Methodology: D.P.M. and J.L.P. Investigation: D.P.M., J.L.P. L.S-W., F.G., and J.B.B. Data analysis: D.P.M. and J.L.P. Software: D.P.M. Visualization: D.P.M. Writing – original draft: - D.P.M. Writing – review and editing: - D.P.M., J.L.P., L.S-W., F.G., and J.B.B.

## Competing interests

The authors have no competing interests to declare.

## Data and material availability

Atomic coordinates for structures reported in this study are deposited in the Protein Data Bank under the accession codes 9D8Q (*rf*ERα LBD bound to E2 and *h*SRC2-2 peptide) and 9D8R (*rf*ERγ LBD bound to E2). Experimental data are deposited on Figshare (DOI: 10.25909/27366339). Code used for analysis and working are deposited online on GitHub (https://github.com/BruningLab).

## Supplementary Materials for

**Fig. S1.**
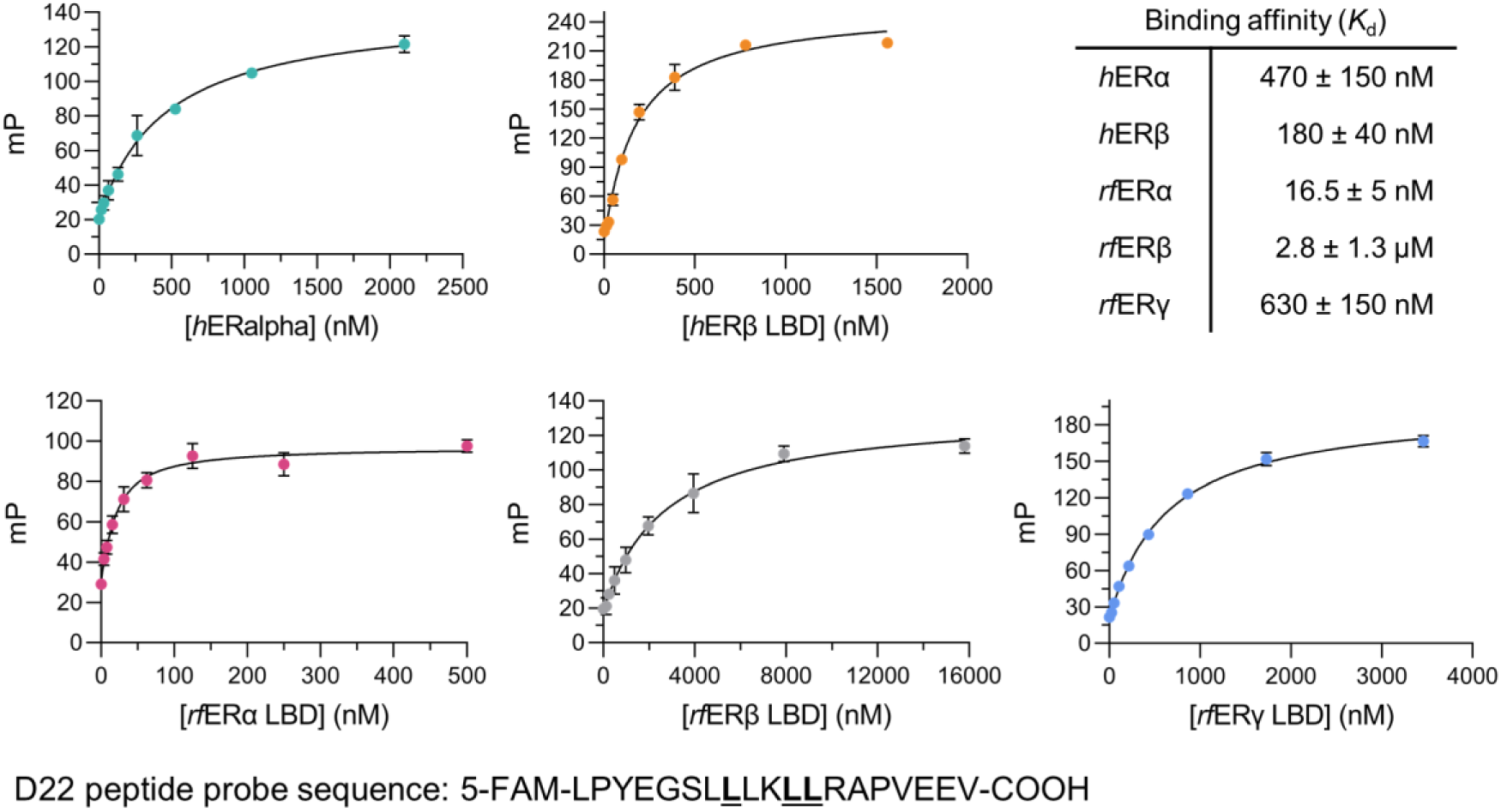
*In vitro* fluorescence polarisation assays demonstrate coactivator recognition using a *LxxLL* peptide probe. The x-axis represents increasing concentrations of recombinant protein (in nanomolar) bound to estradiol (E2) titrated in fluorescence polarisation (FP) buffer containing 10 nM D22 peptide probe. Data represents mean ± standard deviation of three experiments.

**Fig. S2.**
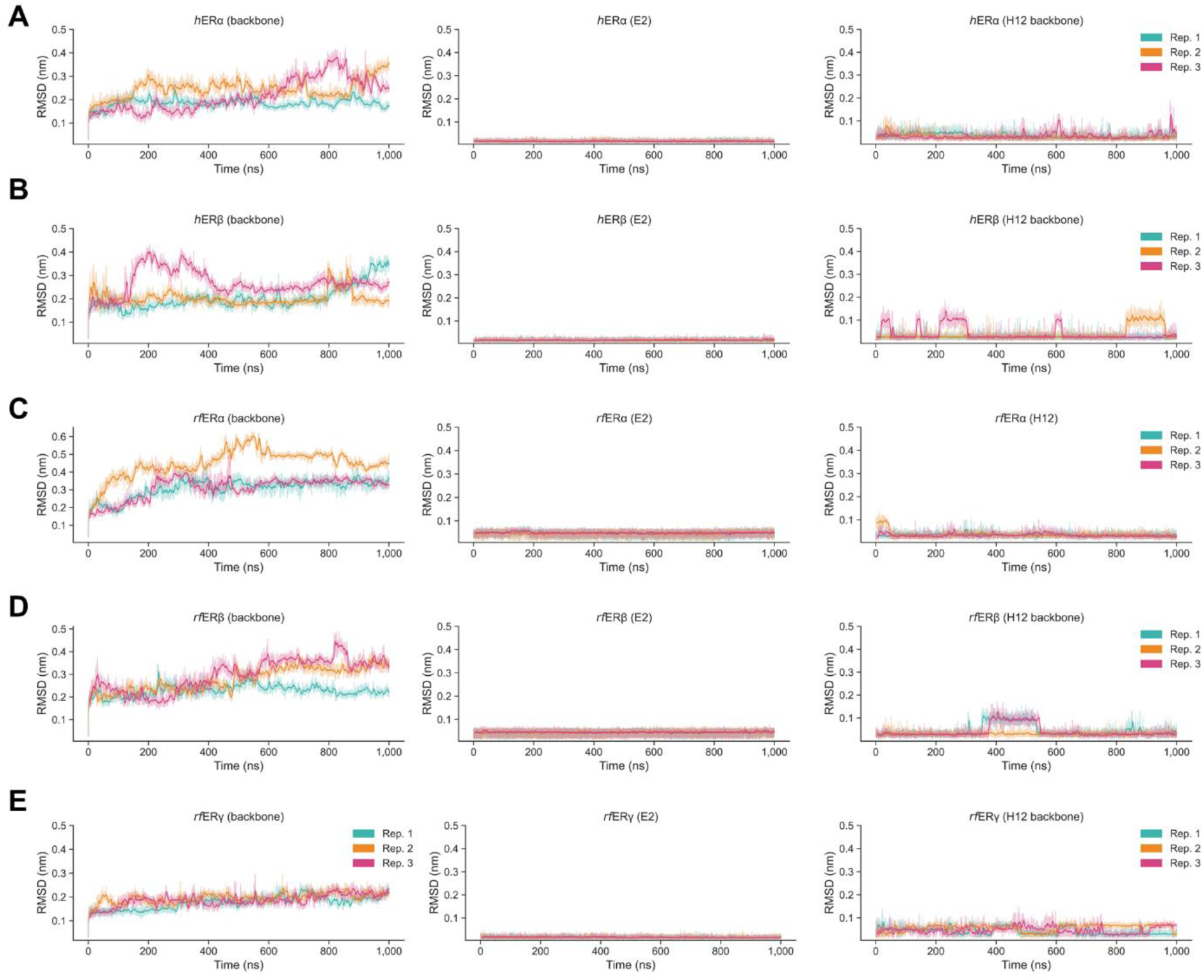
Root mean-squared deviation (RMSD) time-course plots for each homolog. Left: backbone heavy-atom RMSD for each 1 µs replicate; middle: estradiol (E2) RMSD for each 1 µs replicate; right: helix-12 (H12) backbone RMSD for each 1 µs replicate. a-e, in order: *h*ERα LBD, *h*ERβ LBD, *rf*ERα LBD, *rf*ERβ LBD and *rf*ERγ LBD. The 5 ns rolling-average (solid line) is superimposed above raw RMSD values.

**Fig. S3.**
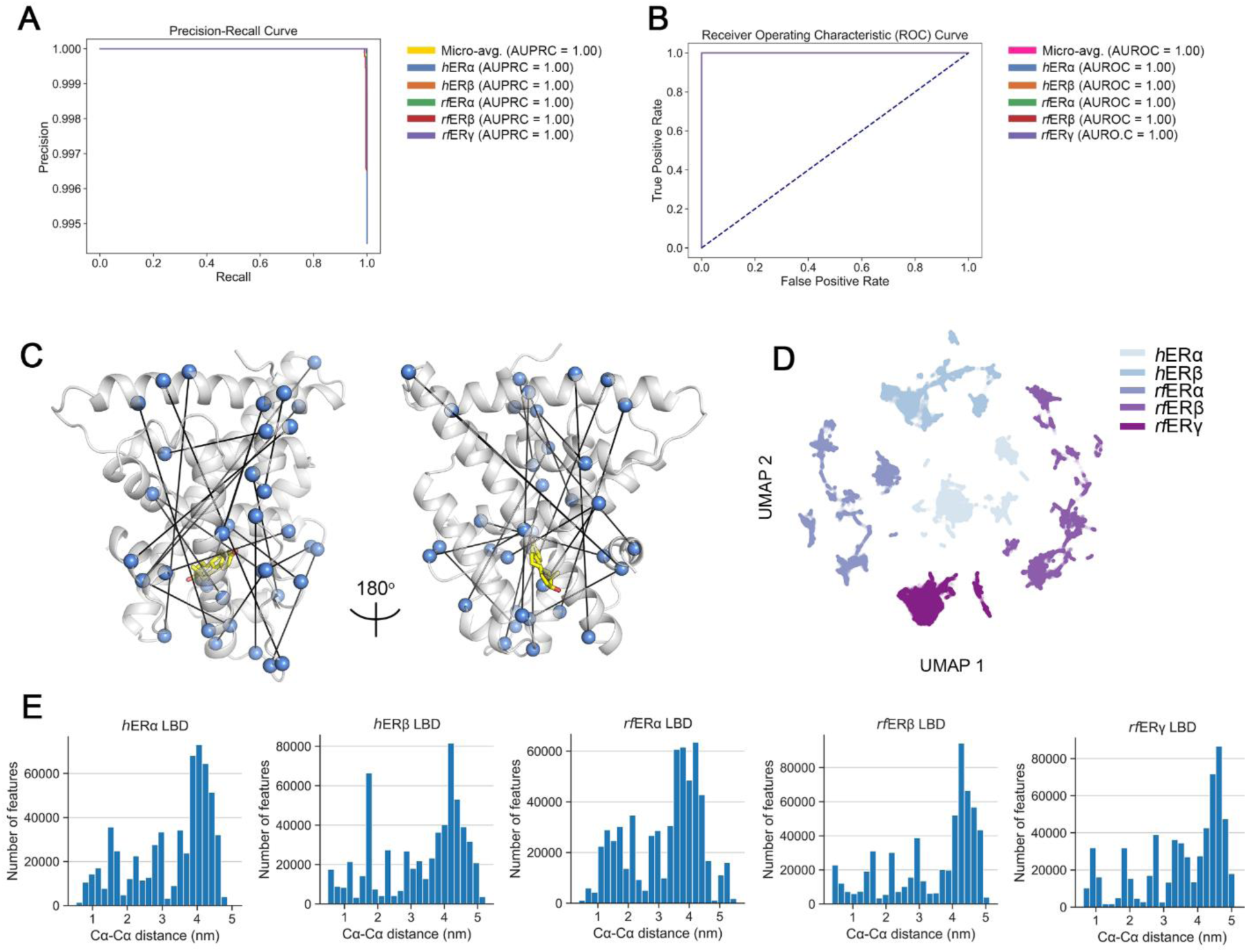
Supervised-learning classification for identification of features used for Markov State modelling. See Methods section “*Markov State modelling: Feature selection”* for model training details. a, Area under the precision-recall curve (AUPRC) and b, area under the receiver operating characteristic curve (AUROC) from evaluation of the hyper-parameter optimised *XGBoost* model on the holdout test set (a randomly shuffled 70:30 train-test split was used). c, Important features (vectors between two Cα-atoms; n=20) were extracted from the model and mapped to structure; shown here is the *rf*ERγ LBD. d, Uniform Manifold Approximation and Projection (UMAP) of the Z-score normalised features identified by the model (n_neighbours = 5, min_dist = 0.2). e, Distance distribution in nm of the features (Cα-Cα) for each homolog extracted from the model.

**Fig. S4.**
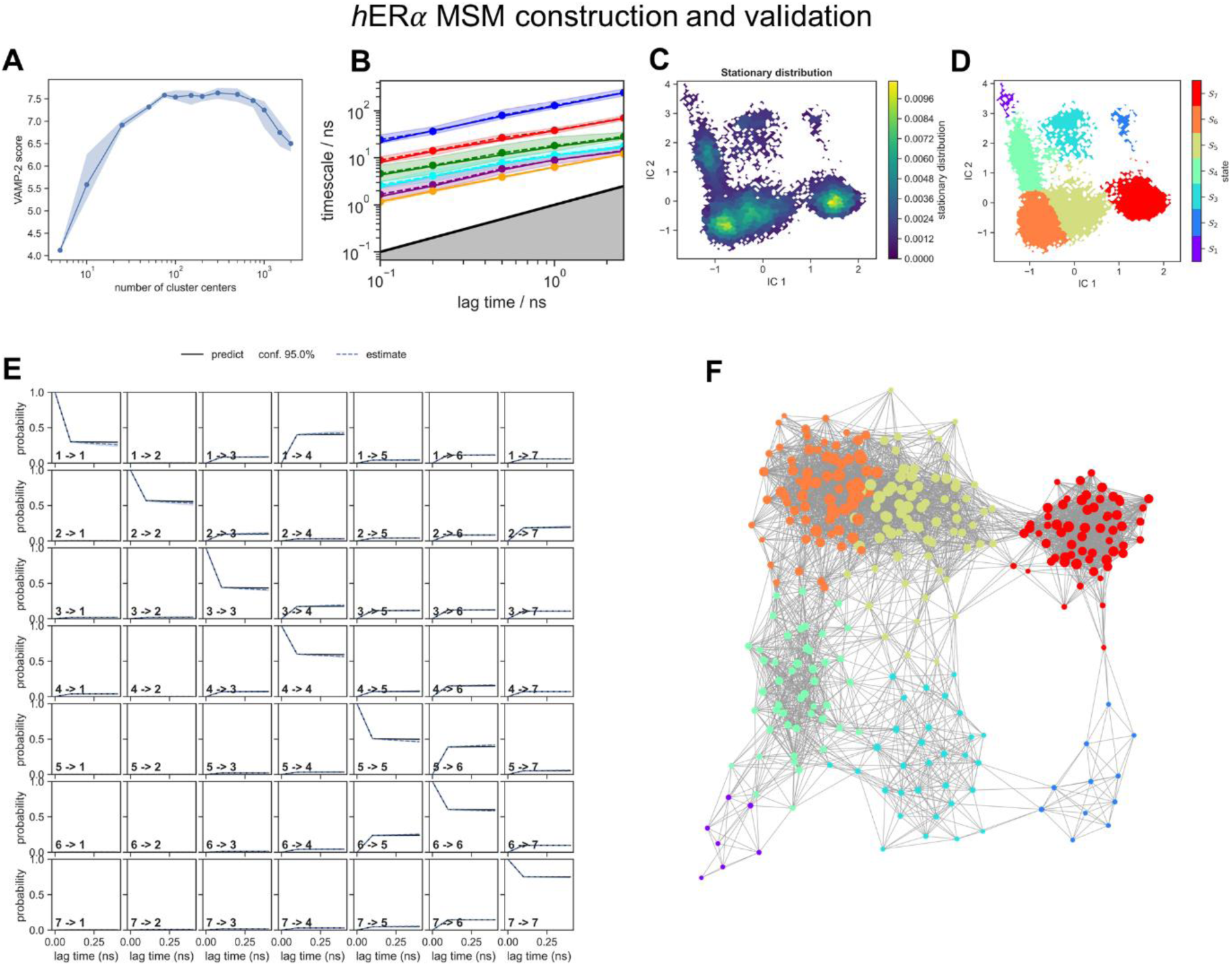
Construction, validation and network visualisation of the *h*ERα LBD Bayesian Markov State modelling. a, Estimation of the optimal number of *k*-means cluster centres (microstates) for Bayesian MSM construction. The blue line represents the mean VAMP-2 score heuristic of five unvalidated MSMs and shading corresponds to the 90% confidence interval. b, Implied timescales (ITS) plot using *k* = 300 microstates and a lag time ***t*** = 1 step (100 ps). c, Stationary distribution ***π*** of discrete microstates from the Bayesian MSM mapped to the first two components of the time-structured independent component analysis (TICA) discretisation of feature space. d, Macrostate assignments determined from coarse-graining the MSM with Perron Cluster Cluster Analysis (PCCA) mapped to feature space. Each colour represents a macrostate assignment. e, Statistical validation of the 7-state Bayesian MSM using the Chapman-Kolmogorov test. f, Network representation of the Bayesian MSM. Each node corresponds to a microstate and node size is proportional to node degree; edges connect nodes if the transition probability is > 0, which is proportional to the edge width. Each microstate is coloured according to macrostate assignment, as shown in panel d. The MSM was constructed in *PyEMMA* using the features identified by the XGBoost supervised-learning classification model. Network visualisation was performed using *CytoScape*.

**Fig. S5.**
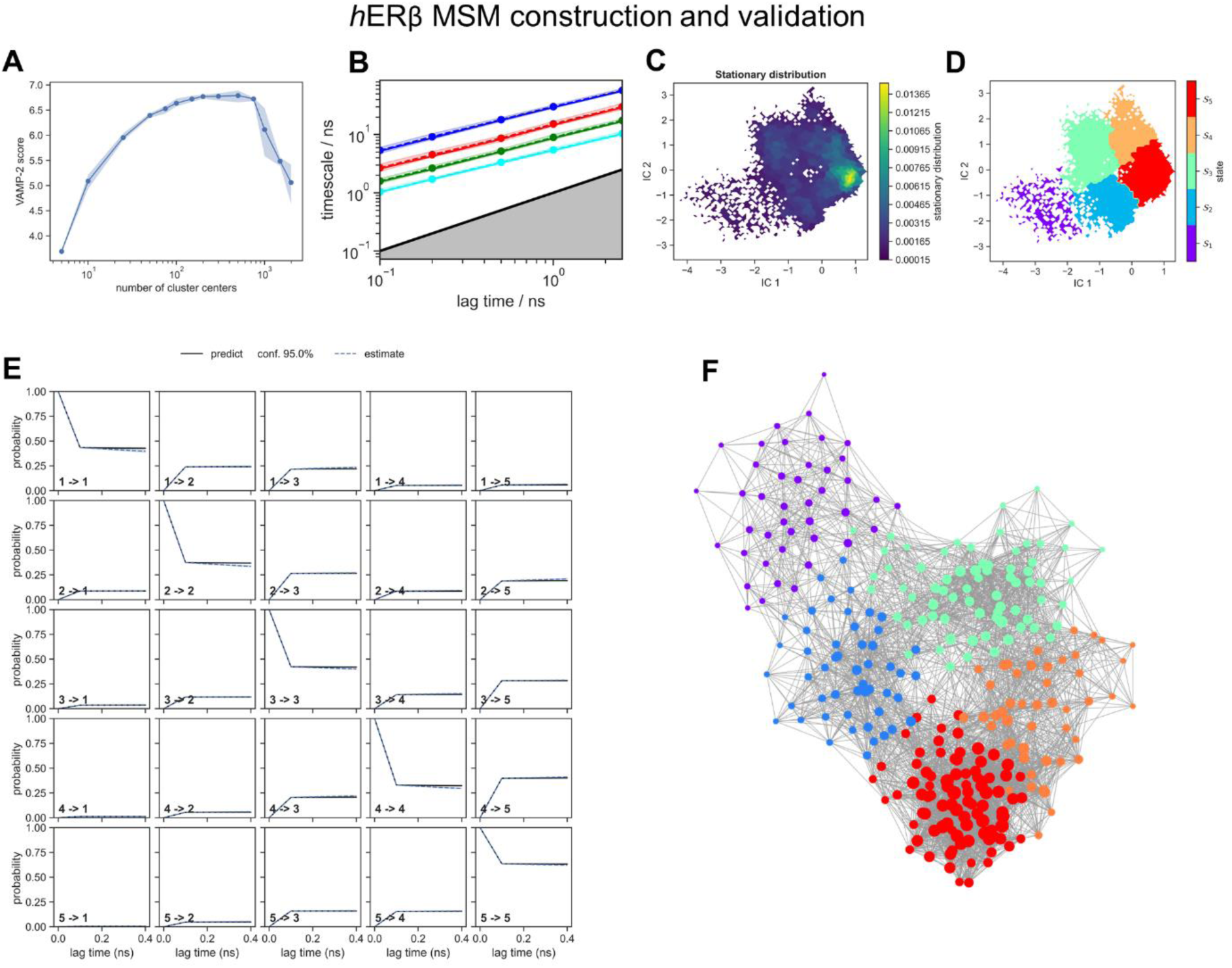
Construction, validation and network visualisation of the *h*ERβ LBD Bayesian Markov State modelling. a, Estimation of the optimal number of *k*-means cluster centres (microstates) for Bayesian MSM construction. The blue line represents the mean VAMP-2 score heuristic of five unvalidated MSMs and shading corresponds to the 90% confidence interval. b, Implied timescales (ITS) plot using *k* = 300 microstates and a lag time ***t*** = 1 step (100 ps). c, Stationary distribution ***π*** of discrete microstates from the Bayesian MSM mapped to the first two components of the time-structured independent component analysis (TICA) discretisation of feature space. d, Macrostate assignments determined from coarse-graining the MSM with Perron Cluster Cluster Analysis (PCCA) mapped to feature space. Each colour represents a macrostate assignment. e, Statistical validation of the 7-state Bayesian MSM using the Chapman-Kolmogorov test. f, Network representation of the Bayesian MSM. Each node corresponds to a microstate and node size is proportional to node degree; edges connect nodes if the transition probability is > 0, which is proportional to the edge width. Each microstate is coloured according to macrostate assignment, as shown in panel d. The MSM was constructed in *PyEMMA* using the features identified by the XGBoost supervised-learning classification model. Network visualisation was performed using *CytoScape*.

**Fig. S6.**
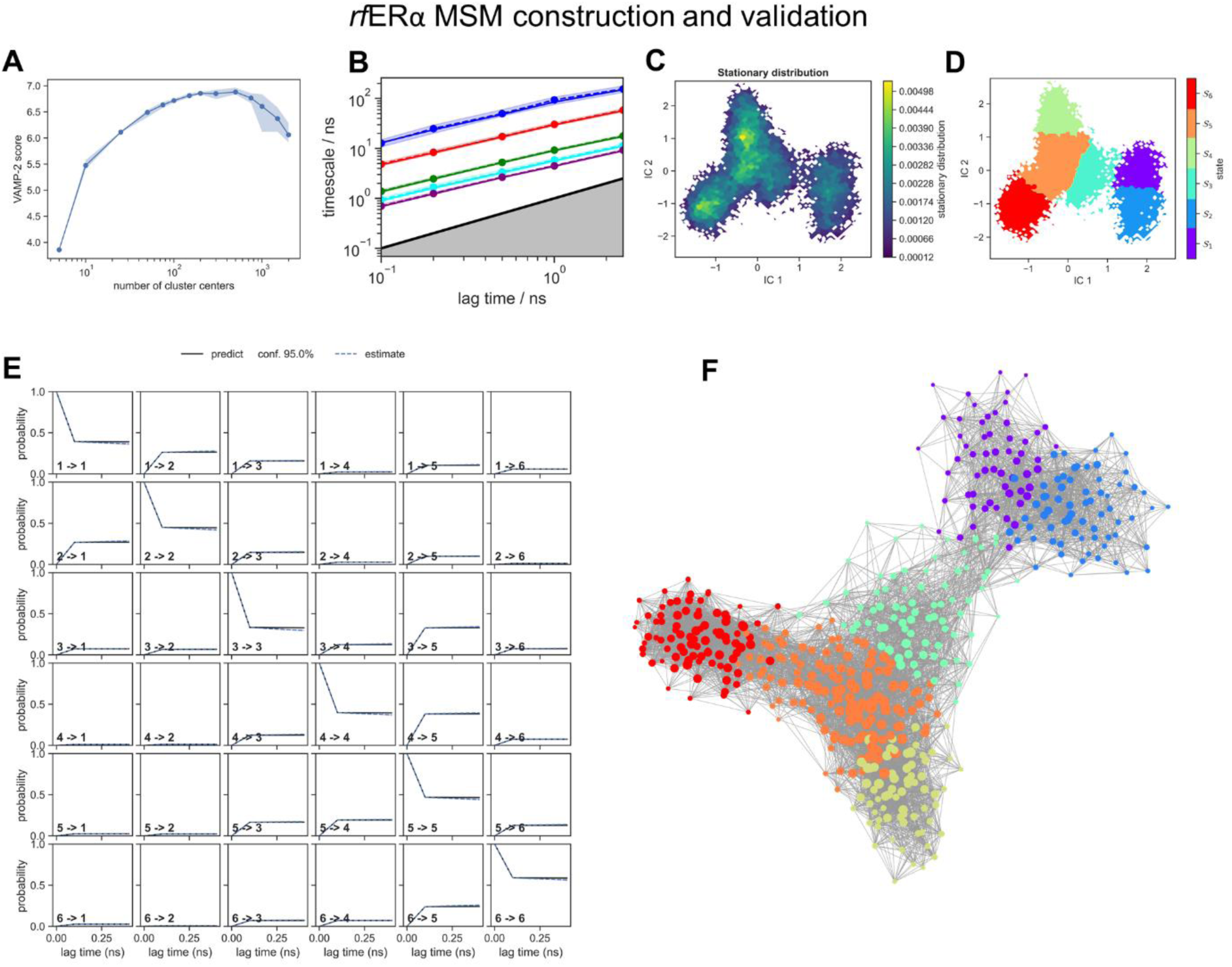
Construction, validation and network visualisation of the *rf*ERα LBD Bayesian Markov State modelling. a, Estimation of the optimal number of *k*-means cluster centres (microstates) for Bayesian MSM construction. The blue line represents the mean VAMP-2 score heuristic of five unvalidated MSMs and shading corresponds to the 90% confidence interval. b, Implied timescales (ITS) plot using *k* = 500 microstates and a lag time ***t*** = 1 step (100 ps). c, Stationary distribution ***π*** of discrete microstates from the Bayesian MSM mapped to the first two components of the time-structured independent component analysis (TICA) discretisation of feature space. d, Macrostate assignments determined from coarse-graining the MSM with Perron Cluster Cluster Analysis (PCCA) mapped to feature space. Each colour represents a macrostate assignment. e, Statistical validation of the 7-state Bayesian MSM using the Chapman-Kolmogorov test. f, Network representation of the Bayesian MSM. Each node corresponds to a microstate and node size is proportional to node degree; edges connect nodes if the transition probability is > 0, which is proportional to the edge width. Each microstate is coloured according to macrostate assignment, as shown in panel d. The MSM was constructed in *PyEMMA* using the features identified by the XGBoost supervised-learning classification model. Network visualisation was performed using *CytoScape*.

**Fig. S7.**
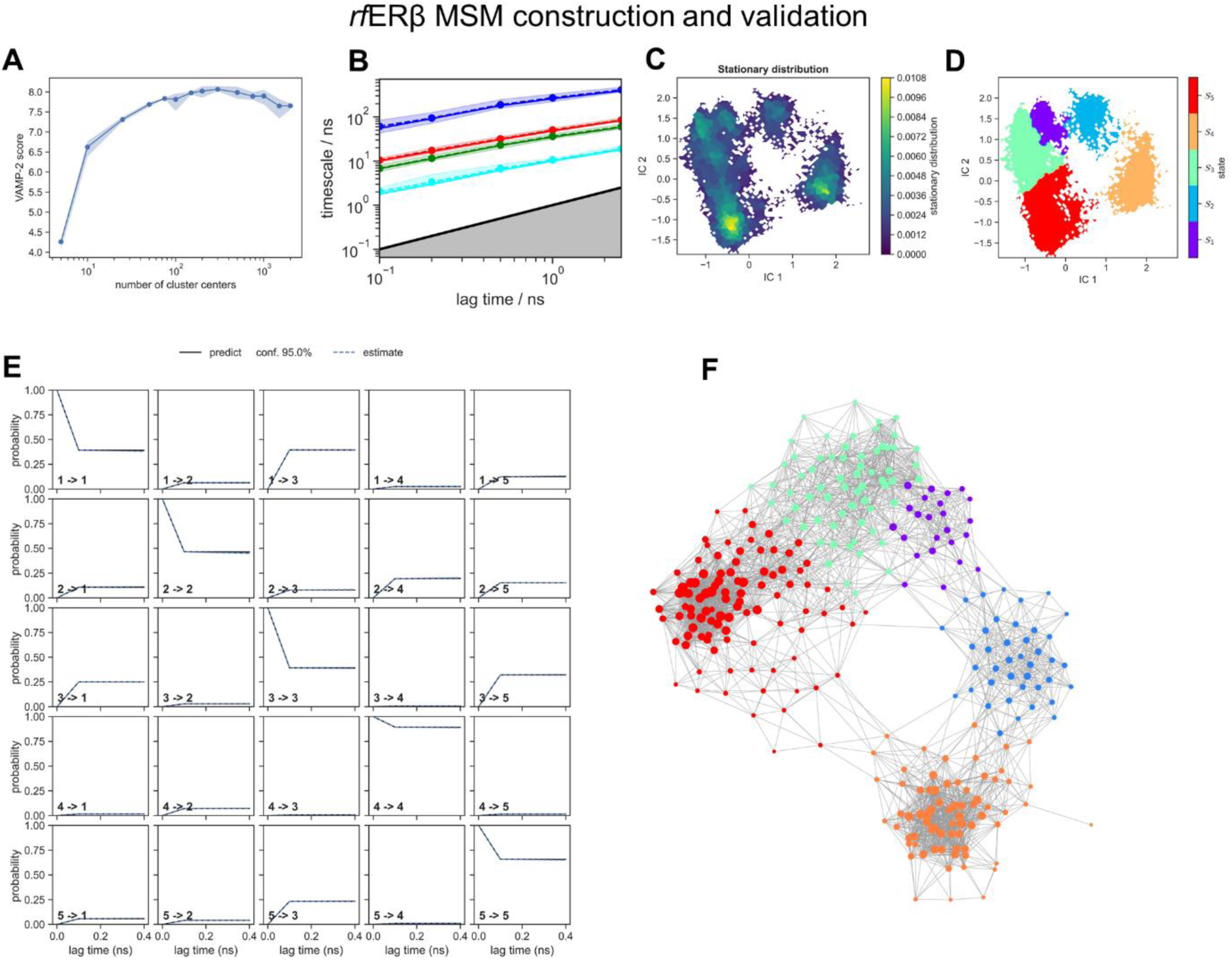
Construction, validation and network visualisation of the *rf*ERβ LBD Bayesian Markov State modelling. **a,** Estimation of the optimal number of *k*-means cluster centres (microstates) for Bayesian MSM construction. The blue line represents the mean VAMP-2 score heuristic of five unvalidated MSMs and shading corresponds to the 90% confidence interval. b, Implied timescales (ITS) plot using *k* = 300 microstates and a lag time ***t*** = 1 step (100 ps). c, Stationary distribution ***π*** of discrete microstates from the Bayesian MSM mapped to the first two components of the time-structured independent component analysis (TICA) discretisation of feature space. d, Macrostate assignments determined from coarse-graining the MSM with Perron Cluster Cluster Analysis (PCCA) mapped to feature space. Each colour represents a macrostate assignment. e, Statistical validation of the 7-state Bayesian MSM using the Chapman-Kolmogorov test. f, Network representation of the Bayesian MSM. Each node corresponds to a microstate and node size is proportional to node degree; edges connect nodes if the transition probability is > 0, which is proportional to the edge width. Each microstate is coloured according to macrostate assignment, as shown in panel d. The MSM was constructed in *PyEMMA* using the features identified by the XGBoost supervised-learning classification model. Network visualisation was performed using *CytoScape*.

**Fig. S8.**
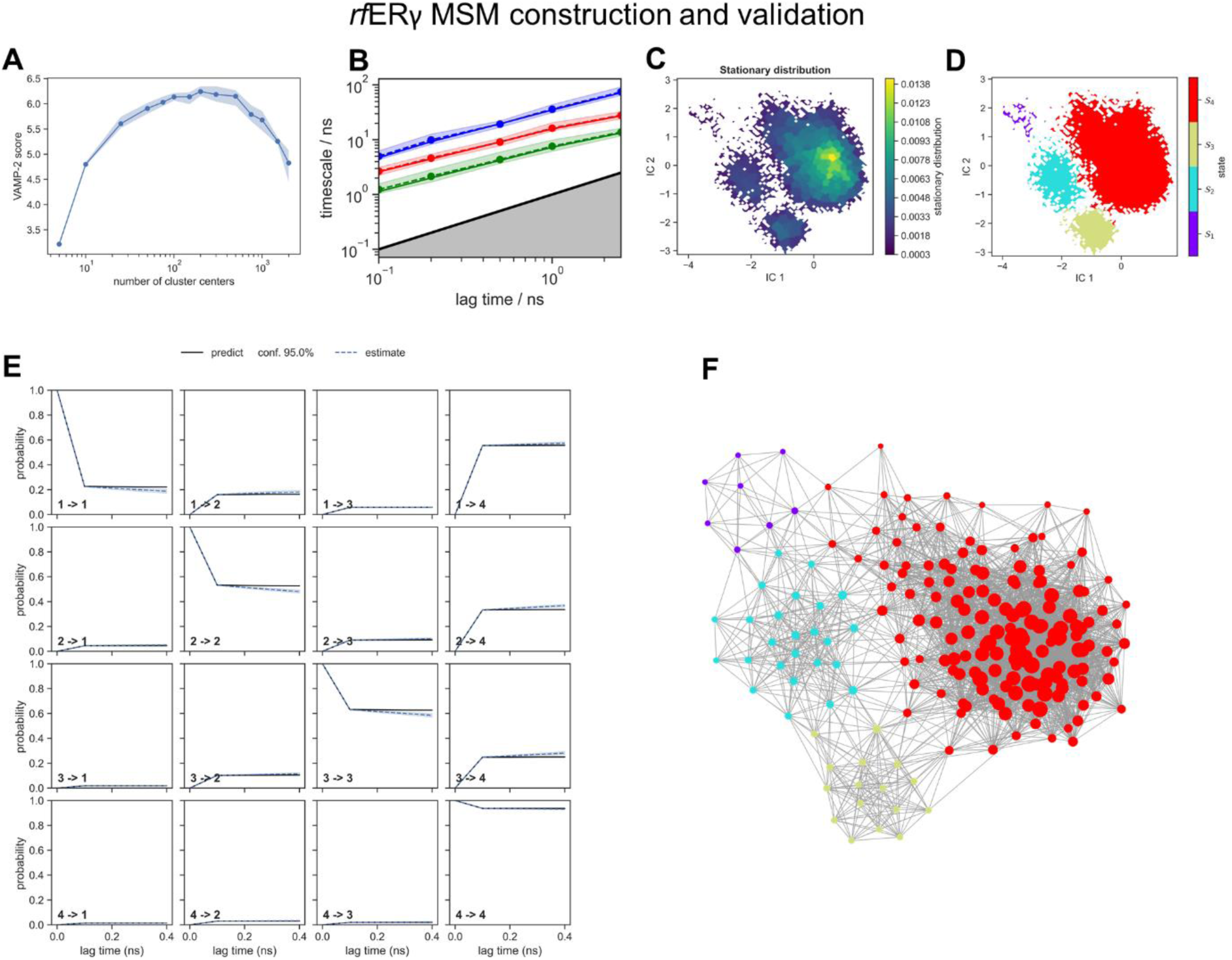
Construction, validation and network visualisation of the *rf*ERγ LBD Bayesian Markov State modelling. a, Estimation of the optimal number of *k*-means cluster centres (microstates) for Bayesian MSM construction. The blue line represents the mean VAMP-2 score heuristic of five unvalidated MSMs and shading corresponds to the 90% confidence interval. b, Implied timescales (ITS) plot using *k* = 200 microstates and a lag time ***t*** = 1 step (100 ps). c, Stationary distribution ***π*** of discrete microstates from the Bayesian MSM mapped to the first two components of the time-structured independent component analysis (TICA) discretisation of feature space. d, Macrostate assignments determined from coarse-graining the MSM with Perron Cluster Cluster Analysis (PCCA) mapped to feature space. Each colour represents a macrostate assignment. e, Statistical validation of the 7-state Bayesian MSM using the Chapman-Kolmogorov test. f, Network representation of the Bayesian MSM. Each node corresponds to a microstate and node size is proportional to node degree; edges connect nodes if the transition probability is > 0, which is proportional to the edge width. Each microstate is coloured according to macrostate assignment, as shown in panel d. The MSM was constructed in *PyEMMA* using the features identified by the XGBoost supervised-learning classification model. Network visualisation was performed using *CytoScape*.

**Fig. S9.**
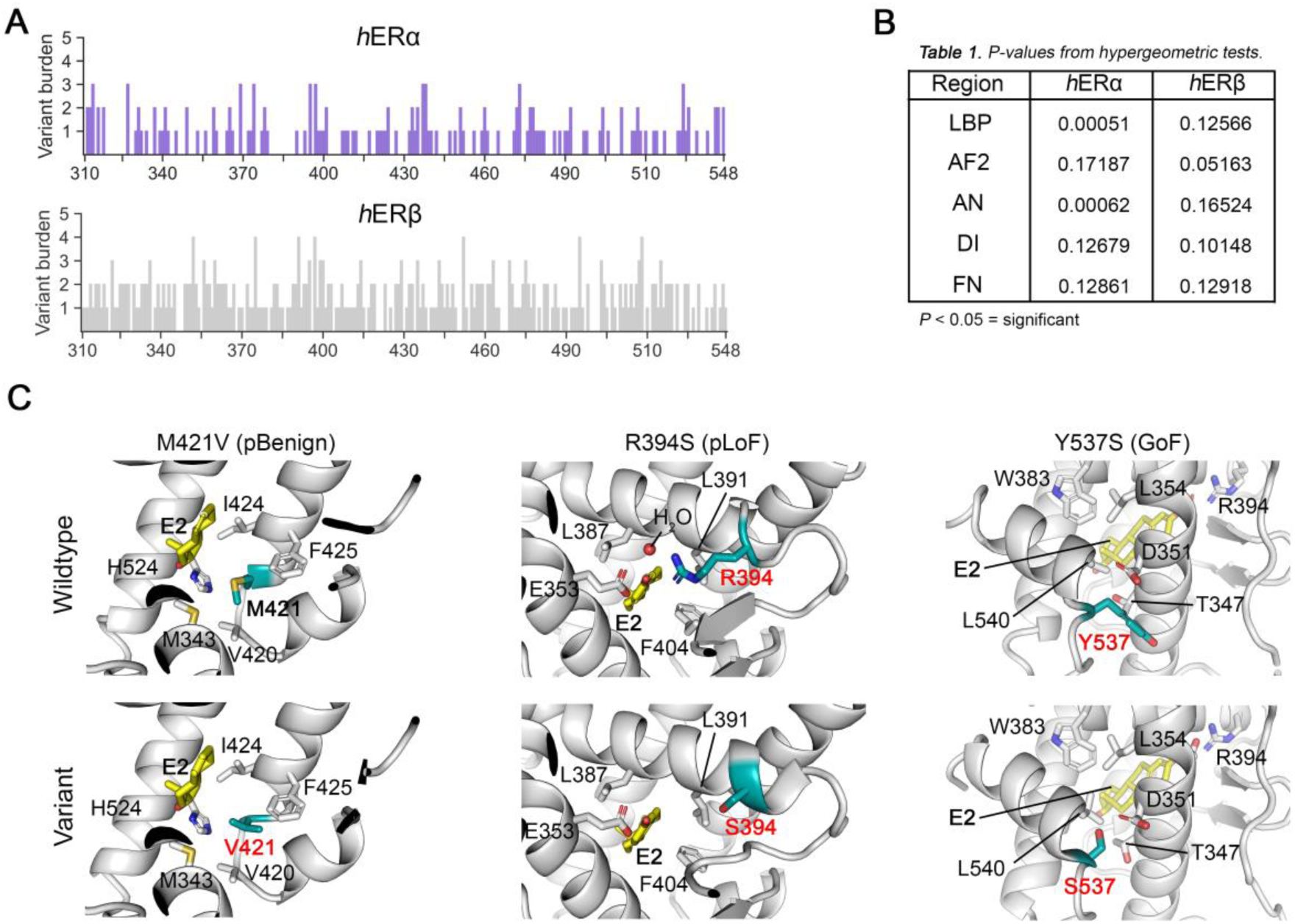
The architecture of genetic variation in hER LBDs. a, Burden of unique missense variants across each position of the *h*ERα and *h*ERβ LBD. Variants were obtained from gnomAD(*48*). b, Significance p-values calculated from hypergeometric tests for statistical depletion of variants in each functional region relative to all positions within the LBD. A significance threshold of ***P*** = 0.05 was used. Only the ligand binding pocket (LBP) and allosteric network (AN) in the *h*ERα LBD were significantly depleted of variants, as compared to the activation function-2 interface (AF2), dimerisation interface (DI) and folding network (NF). No functional region was significantly depleted of variants in the *h*ERβ LBD. c, Structural modelling of example *h*ERα variants M421V (putative benign) and R394S, (putative loss-of-function) and Y537S (gain-of-function; PDB: 3UUD).

**Table S1.**
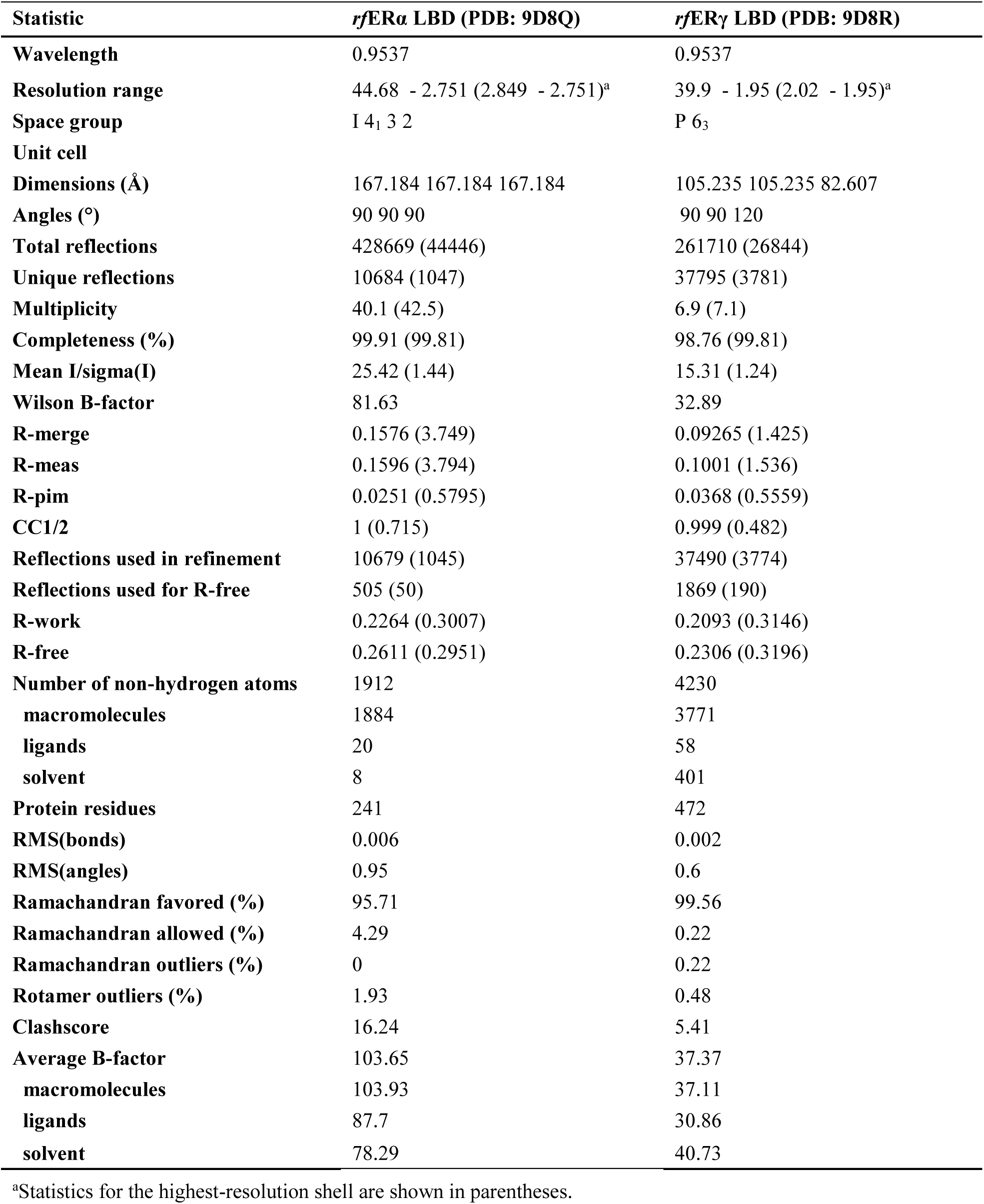
Crystallographic processing and refinement statistics.

| Statistic | *rf*ERα LBD (PDB: 9D8Q) | *rf*ERγ LBD (PDB: 9D8R) |
| --- | --- | --- |
| Wavelength | 0.9537 | 0.9537 |
| Resolution range | 44.68 - 2.751 (2.849 - 2.751)<sup>a</sup> | 39.9 - 1.95 (2.02 - 1.95)<sup>a</sup> |
| Space group | I 4<sub>1</sub> 3 2 | P 6<sub>3</sub> |
| Unit cell |  |  |
| Dimensions (Å) | 167.184 167.184 167.184 | 105.235 105.235 82.607 |
| Angles (°) | 90 90 90 | 90 90 120 |
| Total reflections | 428669 (44446) | 261710 (26844) |
| Unique reflections | 10684 (1047) | 37795 (3781) |
| Multiplicity | 40.1 (42.5) | 6.9 (7.1) |
| Completeness (%) | 99.91 (99.81) | 98.76 (99.81) |
| Mean I/sigma(I) | 25.42 (1.44) | 15.31 (1.24) |
| Wilson B-factor | 81.63 | 32.89 |
| R-merge | 0.1576 (3.749) | 0.09265 (1.425) |
| R-meas | 0.1596 (3.794) | 0.1001 (1.536) |
| R-pim | 0.0251 (0.5795) | 0.0368 (0.5559) |
| CC1/2 | 1 (0.715) | 0.999 (0.482) |
| Reflections used in refinement | 10679 (1045) | 37490 (3774) |
| Reflections used for R-free | 505 (50) | 1869 (190) |
| R-work | 0.2264 (0.3007) | 0.2093 (0.3146) |
| R-free | 0.2611 (0.2951) | 0.2306 (0.3196) |
| Number of non-hydrogen atoms | 1912 | 4230 |
| macromolecules | 1884 | 3771 |
| ligands | 20 | 58 |
| solvent | 8 | 401 |
| Protein residues | 241 | 472 |
| RMS(bonds) | 0.006 | 0.002 |
| RMS(angles) | 0.95 | 0.6 |
| Ramachandran favored (%) | 95.71 | 99.56 |
| Ramachandran allowed (%) | 4.29 | 0.22 |
| Ramachandran outliers (%) | 0 | 0.22 |
| Rotamer outliers (%) | 1.93 | 0.48 |
| Clashscore | 16.24 | 5.41 |
| Average B-factor | 103.65 | 37.37 |
| macromolecules | 103.93 | 37.11 |
| ligands | 87.7 | 30.86 |
| solvent | 78.29 | 40.73 |

<sup>a</sup>Statistics for the highest-resolution shell are shown in parentheses.

## Notes

### Competing Interest Statement

The authors have declared no competing interest.

## References

1 Jia, M., Dahlman-Wright, K. & Gustafsson, J.-Å. Estrogen receptor alpha and beta in health and disease. Best Practice & Research Clinical Endocrinology & Metabolism 29, 557–568, 10.1016/j.beem.2015.04.008 (2015).

2 Mangelsdorf, D. J. et al. The nuclear receptor superfamily: the second decade. Cell 83, 835–839, doi:10.1016/0092-8674(95)90199-x (1995).

3 Chen, P., Li, B. & Ou-Yang, L. Role of estrogen receptors in health and disease. Frontiers in endocrinology 13, 839005, doi:10.3389/fendo.2022.839005 (2022).

4 Walter, P. et al. Cloning of the human estrogen receptor cDNA. Proceedings of the National Academy of Sciences 82, 7889–7893, doi:10.1073/pnas.82.23.7889 (1985).

5 Kuiper, G. G., Enmark, E., Pelto-Huikko, M., Nilsson, S. & Gustafsson, J. A. Cloning of a novel receptor expressed in rat prostate and ovary. Proceedings of the National Academy of Sciences 93, 5925–5930, doi:10.1073/pnas.93.12.5925 (1996).

6 Hawkins Mary, B., et al. Identification of a third distinct estrogen receptor and reclassification of estrogen receptors in teleosts. Proceedings of the National Academy of Sciences 97, 10751–10756, doi:10.1073/pnas.97.20.10751 (2000).

7 Thornton, J. W. Evolution of vertebrate steroid receptors from an ancestral estrogen receptor by ligand exploitation and serial genome expansions. Proceedings of the National Academy of Sciences 98, 5671–5676, doi:10.1073/pnas.091553298 (2001).

8 Tohyama, S. et al. Evolution of estrogen receptors in ray-finned fish and their comparative responses to estrogenic substances. The Journal of Steroid Biochemistry and Molecular Biology 158, 189–197, 10.1016/j.jsbmb.2015.12.009 (2016).

9 Davesne, D. et al. Fossilized cell structures identify an ancient origin for the teleost whole-genome duplication. Proceedings of the National Academy of Sciences 118, e2101780118, doi:10.1073/pnas.2101780118 (2021).

10 Bridgham Jamie, T., Carroll Sean, M. & Thornton Joseph, W. Evolution of Hormone-Receptor Complexity by Molecular Exploitation. Science 312, 97–101, doi:10.1126/science.1123348 (2006).

11 Bridgham, J. T., Ortlund, E. A. & Thornton, J. W. An epistatic ratchet constrains the direction of glucocorticoid receptor evolution. Nature 461, 515–519, doi:10.1038/nature08249 (2009).

12 Bridgham, J. T. et al. Protein Evolution by Molecular Tinkering: Diversification of the Nuclear Receptor Superfamily from a Ligand-Dependent Ancestor. PLOS Biology 8, e1000497, doi:10.1371/journal.pbio.1000497 (2010).

13 Brzozowski, A. M. et al. Molecular basis of agonism and antagonism in the oestrogen receptor. Nature 389, 753–758, doi:10.1038/39645 (1997).

14 Pike, A. C. W. et al. Structure of the ligand-binding domain of oestrogen receptor beta in the presence of a partial agonist and a full antagonist. The EMBO Journal 18, 4608–4618, 10.1093/emboj/18.17.4608 (1999).

15 Valentine, J. E., Kalkhoven, E., White, R., Hoare, S. & Parker, M. G. Mutations in the Estrogen Receptor Ligand Binding Domain Discriminate between Hormone-dependent Transactivation and Transrepression*. Journal of Biological Chemistry 275, 25322–25329, 10.1074/jbc.M002497200 (2000).

16 Wärnmark, A. et al. Interaction of Transcriptional Intermediary Factor 2 Nuclear Receptor Box Peptides with the Coactivator Binding Site of Estrogen Receptor α*. Journal of Biological Chemistry 277, 21862–21868, 10.1074/jbc.M200764200 (2002).

17 Dai Susie, Y., et al. Prediction of the tissue-specificity of selective estrogen receptor modulators by using a single biochemical method. Proceedings of the National Academy of Sciences 105, 7171–7176, doi:10.1073/pnas.0710802105 (2008).

18 Dai, S. Y. et al. Unique ligand binding patterns between estrogen receptor alpha and beta revealed by hydrogen-deuterium exchange. Biochemistry 48, 9668–9676, doi:10.1021/bi901149t (2009).

19 Shiau, A. K. et al. The Structural Basis of Estrogen Receptor/Coactivator Recognition and the Antagonism of This Interaction by Tamoxifen. Cell 95, 927–937, 10.1016/S0092-8674(00)81717-1 (1998).

20 Toy, W. et al. ESR1 ligand-binding domain mutations in hormone-resistant breast cancer. Nature Genetics 45, 1439–1445, doi:10.1038/ng.2822 (2013).

21 Fanning, S. W. et al. Estrogen receptor alpha somatic mutations Y537S and D538G confer breast cancer endocrine resistance by stabilizing the activating function-2 binding conformation. eLife 5, e12792, doi:10.7554/eLife.12792 (2016).

22 Katzenellenbogen, J. A., Mayne, C. G., Katzenellenbogen, B. S., Greene, G. L. & Chandarlapaty, S. Structural underpinnings of oestrogen receptor mutations in endocrine therapy resistance. Nature Reviews Cancer 18, 377–388, doi:10.1038/s41568-018-0001-z (2018).

23 Bernard, V. et al. Familial Multiplicity of Estrogen Insensitivity Associated With a Loss-of-Function ESR1 Mutation. The Journal of clinical endocrinology and metabolism 102, 93–99, doi:10.1210/jc.2016-2749 (2017).

24 Li, Y. et al. A mutant form of ERα associated with estrogen insensitivity affects the coupling between ligand binding and coactivator recruitment. Science Signaling 13, eaaw4653, doi:10.1126/scisignal.aaw4653 (2020).

25 Li, Y. et al. ESR1 Mutations Associated With Estrogen Insensitivity Syndrome Change Conformation of Ligand-Receptor Complex and Altered Transcriptome Profile. Endocrinology 161, bqaa050, doi:10.1210/endocr/bqaa050 (2020).

26 Imamov, O., Shim, G.-J., Warner, M. & Gustafsson, J.-A. k. Estrogen Receptor beta in Health and Disease1. Biology of Reproduction 73, 866–871, doi:10.1095/biolreprod.105.043497 (2005).

27 Bain, P. A., Papanicolaou, A. & Kumar, A. Identification of Putative Nuclear Receptors and Steroidogenic Enzymes in Murray-Darling Rainbowfish (Melanotaenia fluviatilis) Using RNA-Seq and De Novo Transcriptome Assembly. PLOS ONE 10, e0142636, doi:10.1371/journal.pone.0142636 (2015).

28 Kumar, S. et al. TimeTree 5: An Expanded Resource for Species Divergence Times. Molecular Biology and Evolution 39, msac174, doi:10.1093/molbev/msac174 (2022).

29 Menuet, A. et al. Molecular Characterization of Three Estrogen Receptor Forms in Zebrafish: Binding Characteristics, Transactivation Properties, and Tissue Distributions1. Biology of Reproduction 66, 1881–1892, doi:10.1095/biolreprod66.6.1881 (2002).

30 Tohyama, S. et al. Understanding the Molecular Basis for Differences in Responses of Fish Estrogen Receptor Subtypes to Environmental Estrogens. Environmental Science & Technology 49, 7439–7447, doi:10.1021/acs.est.5b00704 (2015).

31 Ekena, K., Weis, K. E., Katzenellenbogen, J. A. & Katzenellenbogen, B. S. Identification of Amino Acids in the Hormone Binding Domain of the Human Estrogen Receptor Important in Estrogen Binding*. Journal of Biological Chemistry 271, 20053–20059, 10.1074/jbc.271.33.20053(1996).

32 Hawkins, M. B. & Thomas, P. The Unusual Binding Properties of the Third Distinct Teleost Estrogen Receptor Subtype ERβa Are Accompanied by Highly Conserved Amino Acid Changes in the Ligand Binding Domain. Endocrinology 145, 2968–2977, doi:10.1210/en.2003-0806 (2004).

33 Ekena, K., Weis, K. E., Katzenellenbogen, J. A. & Katzenellenbogen, B. S. Identification of amino acids in the hormone binding domain of the human estrogen receptor important in estrogen binding. The Journal of biological chemistry 271, 20053–20059, doi:10.1074/jbc.271.33.20053 (1996).

34 Chothia, C. & Janin, J. Principles of protein–protein recognition. Nature 256, 705–708, doi:10.1038/256705a0 (1975).

35 Conte, L. L., Chothia, C. & Janin, J. The atomic structure of protein-protein recognition sites11Edited by A. R. Fersht. Journal of Molecular Biology 285, 2177–2198, 10.1006/jmbi.1998.2439 (1999).

36 Podgornaia, A. I. & Laub, M. T. Pervasive degeneracy and epistasis in a protein-protein interface. Science 347, 673–677, doi:10.1126/science.1257360(2015).

37 Atwell, S., Ultsch, M., De Vos, A. M. & Wells, J. A. Structural Plasticity in a Remodeled Protein-Protein Interface. Science 278, 1125–1128, doi:10.1126/science.278.5340.1125 (1997).

38 Starr, T. N. & Thornton, J. W. Epistasis in protein evolution. Protein Sci 25, 1204–1218, doi:10.1002/pro.2897 (2016).

39 Hochberg, G. K. A. et al. A hydrophobic ratchet entrenches molecular complexes. Nature 588, 503–508, doi:10.1038/s41586-020-3021-2 (2020).

40 Noé, F., Schütte, C., Vanden-Eijnden, E., Reich, L. & Weikl, T. R. Constructing the equilibrium ensemble of folding pathways from short off-equilibrium simulations. Proceedings of the National Academy of Sciences 106, 19011–19016, doi:10.1073/pnas.0905466106 (2009).

41 Prinz, J.-H. et al. Markov models of molecular kinetics: Generation and validation. The Journal of Chemical Physics 134, 174105, doi:10.1063/1.3565032 (2011).

42 Wrenn, C. K. & Katzenellenbogen, B. S. Structure-function analysis of the hormone binding domain of the human estrogen receptor by region-specific mutagenesis and phenotypic screening in yeast. Journal of Biological Chemistry 268, 24089–24098, 10.1016/S0021-9258(20)80497-9 (1993).

43 Herynk, M. H. & Fuqua, S. A. W. Estrogen Receptor Mutations in Human Disease. Endocrine Reviews 25, 869–898, doi:10.1210/er.2003-0010 (2004).

44 Reese, J. C. & Katzenellenbogen, B. S. Mutagenesis of cysteines in the hormone binding domain of the human estrogen receptor. Alterations in binding and transcriptional activation by covalently and reversibly attaching ligands. Journal of Biological Chemistry 266, 10880–10887, 10.1016/S0021-9258(18)99101-5 (1991).

45 Thornton Joseph, W., Need, E. & Crews, D. Resurrecting the Ancestral Steroid Receptor: Ancient Origin of Estrogen Signaling. Science 301, 1714–1717, doi:10.1126/science.1086185 (2003).

46 Keay, J., Bridgham, J. T. & Thornton, J. W. The Octopus vulgaris Estrogen Receptor Is a Constitutive Transcriptional Activator: Evolutionary and Functional Implications. Endocrinology 147, 3861–3869, doi:10.1210/en.2006-0363 (2006).

47 Bridgham, J. T., Keay, J., Ortlund, E. A. & Thornton, J. W. Vestigialization of an Allosteric Switch: Genetic and Structural Mechanisms for the Evolution of Constitutive Activity in a Steroid Hormone Receptor. PLOS Genetics 10, e1004058, doi:10.1371/journal.pgen.1004058 (2014).

48 Karczewski, K. J. et al. The mutational constraint spectrum quantified from variation in 141,456 humans. Nature 581, 434–443, doi:10.1038/s41586-020-2308-7 (2020).

49 Gudmundsson, S. et al. Variant interpretation using population databases: Lessons from gnomAD. Human mutation 43, 1012–1030, doi:10.1002/humu.24309 (2022).

50 Sun, B. B. et al. Genetic associations of protein-coding variants in human disease. Nature 603, 95–102, doi:10.1038/s41586-022-04394-w (2022).

51 MacArthur, D. G. et al. Guidelines for investigating causality of sequence variants in human disease. Nature 508, 469–476, doi:10.1038/nature13127 (2014).

52 Nasrazadani, A., Thomas, R. A., Oesterreich, S. & Lee, A. V. Precision Medicine in Hormone Receptor-Positive Breast Cancer. Frontiers in Oncology 8, doi:10.3389/fonc.2018.00144 (2018).

53 Nettles, K. W. et al. NFκB selectivity of estrogen receptor ligands revealed by comparative crystallographic analyses. Nature Chemical Biology 4, 241–247, doi:10.1038/nchembio.76 (2008).

54 Ozers, M. S. et al. Analysis of Ligand-Dependent Recruitment of Coactivator Peptides to Estrogen Receptor Using Fluorescence Polarization. Molecular Endocrinology 19, 25–34, doi:10.1210/me.2004-0256 (2005).

55 Pederick, J. L., Frkic, R. L., McDougal, D. P. & Bruning, J. B. A structural basis for the activation of peroxisome proliferator-activated receptor gamma (PPARγ) by perfluorooctanoic acid (PFOA). Chemosphere 354, 141723, 10.1016/j.chemosphere.2024.141723 (2024).

56 Abagyan, R. et al. Homology modeling with internal coordinate mechanics: Deformation zone mapping and improvements of models via conformational search. Proteins: Structure, Function, and Bioinformatics 29, 29–37, 10.1002/(SICI)1097-0134(1997)1+<29::AID-PROT5>3.0.CO;2-J (1997).

57 Cowieson, N. P. et al. MX1: a bending-magnet crystallography beamline serving both chemical and macromolecular crystallography communities at the Australian Synchrotron. Journal of Synchrotron Radiation 22, 187–190, doi:doi:10.1107/S1600577514021717 (2015).

58 Aragao, D. et al. MX2: a high-flux undulator microfocus beamline serving both the chemical and macromolecular crystallography communities at the Australian Synchrotron. Journal of Synchrotron Radiation 25, 885–891, doi:doi:10.1107/S1600577518003120 (2018).

59 Kabsch, W. XDS. Acta Crystallographica Section D 66, 125–132, doi:doi:10.1107/S0907444909047337 (2010).

60 Winn, M. D. et al. Overview of the CCP4 suite and current developments. Acta Crystallographica Section D 67, 235–242, doi:doi:10.1107/S0907444910045749 (2011).

61 Emsley, P., Lohkamp, B., Scott, W. G. & Cowtan, K. Features and development of Coot. *A*cta Crystallogr D Biol Crystallogr 66, 486–501, doi:10.1107/s0907444910007493 (2010).

62 Liebschner, D. et al. Macromolecular structure determination using X-rays, neutrons and electrons: recent developments in Phenix. Acta Crystallogr D Struct Biol 75, 861–877, doi:10.1107/s2059798319011471 (2019).

63 McCoy, A. J. et al. Phaser crystallographic software. Journal of Applied Crystallography 40, 658–674, doi:doi:10.1107/S0021889807021206 (2007).

64 Stein, N. CHAINSAW: a program for mutating pdb files used as templates in molecular replacement. Journal of Applied Crystallography 41, 641–643, doi:doi:10.1107/S0021889808006985 (2008).

65 McGibbon, R. T. et al. MDTraj: A Modern Open Library for the Analysis of Molecular Dynamics Trajectories. Biophys J 109, 1528–1532, doi:10.1016/j.bpj.2015.08.015 (2015).

66 Harris, C. R. et al. Array programming with NumPy. Nature 585, 357–362, doi:10.1038/s41586-020-2649-2 (2020).

67 Shannon, P. et al. Cytoscape: a software environment for integrated models of biomolecular interaction networks. Genome Res 13, 2498–2504, doi:10.1101/gr.1239303 (2003).

68 Van Der Spoel, D. et al. GROMACS: Fast, flexible, and free. J Comput Chem 26, 1701–1718, 10.1002/jcc.20291 (2005).

69 Abraham, M. J. et al. GROMACS: High performance molecular simulations through multi-level parallelism from laptops to supercomputers. SoftwareX 1-2, 19–25, 10.1016/j.softx.2015.06.001 (2015).

70 Vanommeslaeghe, K. et al. CHARMM general force field: A force field for drug-like molecules compatible with the CHARMM all-atom additive biological force fields. J Comput Chem 31, 671–690, 10.1002/jcc.21367 (2010).

71 Huang, J. et al. CHARMM36m: an improved force field for folded and intrinsically disordered proteins. Nat Methods 14, 71–73, doi:10.1038/nmeth.4067 (2017).

72 Zoete, V., Cuendet, M. A., Grosdidier, A. & Michielin, O. SwissParam: A fast force field generation tool for small organic molecules. J Comput Chem 32, 2359–2368, 10.1002/jcc.21816 (2011).

73 Hunter, J. D. Matplotlib: A 2D Graphics Environment. Computing in Science & Engineering 9, 90–95, doi:10.1109/MCSE.2007.55 (2007).

74 Waskom, M. L. Seaborn: statistical data visualization. Journal of Open Source Software 6, 3021 (2021).

75 Dai, S. Y. et al. Unique Ligand Binding Patterns between Estrogen Receptor α and β Revealed by Hydrogen−Deuterium Exchange. Biochemistry 48, 9668–9676, doi:10.1021/bi901149t (2009).

76 Pedregosa, F. et al. Scikit-learn: Machine learning in Python. the Journal of machine Learning research 12, 2825–2830 (2011).

77 Chen, T. & Guestrin, C. in Proceedings of the 22nd ACM SIGKDD International Conference on Knowledge Discovery and Data Mining 785– 794 (Association for Computing Machinery, San Francisco, California, USA, 2016).

78 McInnes, L., Healy, J. & Melville, J. Umap: Uniform manifold approximation and projection for dimension reduction. *arXiv preprint arXiv:1802.03426* (2018).

79 Scherer, M. K. et al. PyEMMA 2: A Software Package for Estimation, Validation, and Analysis of Markov Models. Journal of Chemical Theory and Computation 11, 5525–5542, doi:10.1021/acs.jctc.5b00743 (2015).

80 Hagberg, A., Swart, P. & S Chult, D. Exploring network structure, dynamics, and function using NetworkX. (Los Alamos National Lab.(LANL), Los Alamos, NM (United States), 2008).

81 Sayers, E. W. et al. Database resources of the national center for biotechnology information. Nucleic Acids Res 50, D20–d26, doi:10.1093/nar/gkab1112 (2022).

82 Altschul, S. F. et al. Gapped BLAST and PSI-BLAST: a new generation of protein database search programs. Nucleic Acids Res 25, 3389–3402, doi:10.1093/nar/25.17.3389 (1997).

83 Katoh, K. & Standley, D. M. MAFFT multiple sequence alignment software version 7: improvements in performance and usability. Mol Biol Evol 30, 772–780, doi:10.1093/molbev/mst010 (2013).

84 Hoang, D. T., Chernomor, O., von Haeseler, A., Minh, B. Q. & Vinh, L. S. UFBoot2: Improving the Ultrafast Bootstrap Approximation. Molecular Biology and Evolution 35, 518–522, doi:10.1093/molbev/msx281 (2018).

85 Minh, B. Q. et al. IQ-TREE 2: New Models and Efficient Methods for Phylogenetic Inference in the Genomic Era. Molecular Biology and Evolution 37, 1530–1534, doi:10.1093/molbev/msaa015 (2020).

86 Kalyaanamoorthy, S., Minh, B. Q., Wong, T. K. F., von Haeseler, A. & Jermiin, L. S. ModelFinder: fast model selection for accurate phylogenetic estimates. Nature Methods 14, 587–589, doi:10.1038/nmeth.4285 (2017).

87 Capra, J. A. & Singh, M. Predicting functionally important residues from sequence conservation. Bioinformatics 23, 1875–1882, doi:10.1093/bioinformatics/btm270 (2007).

88 Dijkstra, E. W. A note on two problems in connexion with graphs. Numerische Mathematik 1, 269–271 (1959).

89 Shapovalov, Maxim V. & Dunbrack, Roland L. A Smoothed Backbone-Dependent Rotamer Library for Proteins Derived from Adaptive Kernel Density Estimates and Regressions. Structure 19, 844–858, 10.1016/j.str.2011.03.019 (2011).

90 Kabsch, W. & Sander, C. Dictionary of protein secondary structure: Pattern recognition of hydrogen-bonded and geometrical features. Biopolymers 22, 2577–2637, 10.1002/bip.360221211 (1983).

